# An effective method of measuring nanobody binding kinetics and competition-based epitope mapping using biolayer interferometry

**DOI:** 10.1101/2024.10.03.616354

**Authors:** Timothy A. Bates, Sintayehu K. Gurmesa, Jules B. Weinstein, Mila Trank-Greene, Xammy Huu Wrynla, Aidan Anastas, Teketay Wassie Anley, Audrey Hinchliff, Ujwal Shinde, John E. Burke, Fikadu G. Tafesse

## Abstract

Protein-protein interactions (PPI) underpin nearly all biological processes, and understanding the molecular mechanisms governing these interactions is crucial for the progress of biomedical sciences. The emergence of AI-driven computational tools can help reshape the methods in structural biology, however model data often quires empirical validation. The large scale of predictive modeling data will therefore benefit from optimized methodologies for the high-throughput biochemical characterization of PPIs. Biolayer interferometry (BLI) is one of very few approaches that can determine the rate of biomolecular interactions, called kinetics, and of the commonly available kinetic measurement techniques, it is the most suitable for high-throughput experimental designs. Here, we provide step-by-step instructions on how to perform kinetics experiments using BLI. We further describe the basis and execution of competition and epitope binning experiments, which are particularly useful for antibody and nanobody screening applications. The procedure requires 3 hours to complete and is suitable for users with minimal experience with biochemical techniques.

## Introduction

Quantifying the strength of biomolecular interactions is a cornerstone of biochemistry, and a critical step in the development of new medicines. Understanding the binding strength of proteins, such as nanobodies, provides an objective measure of quality and is used in drug discovery campaigns to find leads, in optimization experiments to quantify progress, and in quality control to ensure quality and purity. There are numerous ways to measure binding strength, but Biolayer Interferometry (BLI) is one of very few that can directly measure the rate of binding, also called kinetics ^1^. This is particularly important during discovery and optimization of high-affinity binders because it is essential to differentiate between fast and slow binders which can appear to have similar affinity in simpler binding assays.

This is particularly useful for nanobodies (also called VHHs or single-domain antibodies), which represent a major up-and-coming technology to compete with monoclonal antibodies ^2^. Derived from heavy-chain-only antibodies from camelids or sharks, nanobodies have exceptional properties that make them particularly good candidates for development of affinity reagents and molecular medicines ^3^. Their small size (12-17 kDa) relative to traditional monoclonal antibodies (150 kDa) and single-chain nature are a key feature for their functional utility, overall stability, and speed of development. Because of their simplicity, several bottlenecks in traditional antibody discovery are relieved, particularly the throughput of early discovery processes such as constructing the initial candidate library. Using the most advanced current methods like yeast-, phage-, and ribosome-display, a single researcher can screen over 100 trillion nanobody candidates in a single benchtop experiment. Along with advancements in sequencing technology, it becomes feasible to identify and produce thousands of lead candidates in relatively short timeframes ^4–7^. A new bottleneck then arises at the stage of testing screening hits, both for selection of top candidates, and for improving selection tools. BLI is a powerful tool for the characterization of nanobody candidates including binding kinetics and epitope determination ^8–12^, and can also be used for rapid testing of existing antibodies ^13^.

The same approach can be applied to answer a spectrum of biological questions about the protein complex under study. Affinities of different components of multi-protein complexes can be determined, individual components can be perturbed using nanobodies, and point mutations can be used to validate binding interfaces ^9,10,14,15^. The empirical data provided by these experiments can also be useful for validation of computational models. The success of existing protein folding models can be attributed, in large part, to the wealth of high-quality, annotated structural data in databases like the Protein Data Bank ^16,17^. Databases like BioGRID and IntAct exist for compiling data about PPIs, and expanding databases like these with more experimentally validated affinity data should help provide the critical mass of PPI training data necessary to construct robust computational models ^18,19^.

BLI is a cost-effective and rapid method to measure binding kinetics, but this requires some understanding of how the technique works from a practical perspective as well as a basic framework for results interpretation and troubleshooting. The objective of this work is to provide an easy to use and relatively comprehensive guide to designing, performing, and analyzing nanobody kinetic binding with BLI.

### The Principle of Biolayer Interferometry

BLI is an optical technique which utilizes interferometry to precisely measure the thickness of a biomolecular layer attached to a biosensor surface. It is based on the observation that light can be reflected off a thin layer of protein attached to the end of a small piece of fiber-optic cable. By adding or removing protein to the sensor, the observed length of the fiber increases or decreases by a measurable amount, with this change being proportional to the change in mass. This fine precision is made possible by using an optical setup called an interferometer to measure the pattern of constructive and destructive interference between two parallel beams of light. In this case, the specific layout of components is called a Fabry-Perot interferometer, and it is composed of two parallel reflective surfaces, one at a fixed length from the detector, and one at a variable length (**Figure 1A**) ^20–22^. When the sensor is empty the light returns from fixed point 1 (a thin layer of tantalum oxide), and also from the chemically modified surface of the biosensor at point 2. These light beams combine to give a pattern of different intensities based on the wavelength which serves as the baseline signal (**Figure 1B**). When another molecule is bound to the surface of the sensor, point 2 is no longer reflective because the refractive index now changes a small distance further at point 3 where the new molecule sits. This change in physical position of the reflective plane causes the interference pattern to shift, and the wavelength shift of this change can be plotted over time as more and more molecules bind to the surface (**Figure 1C**).

**Figure 1.**
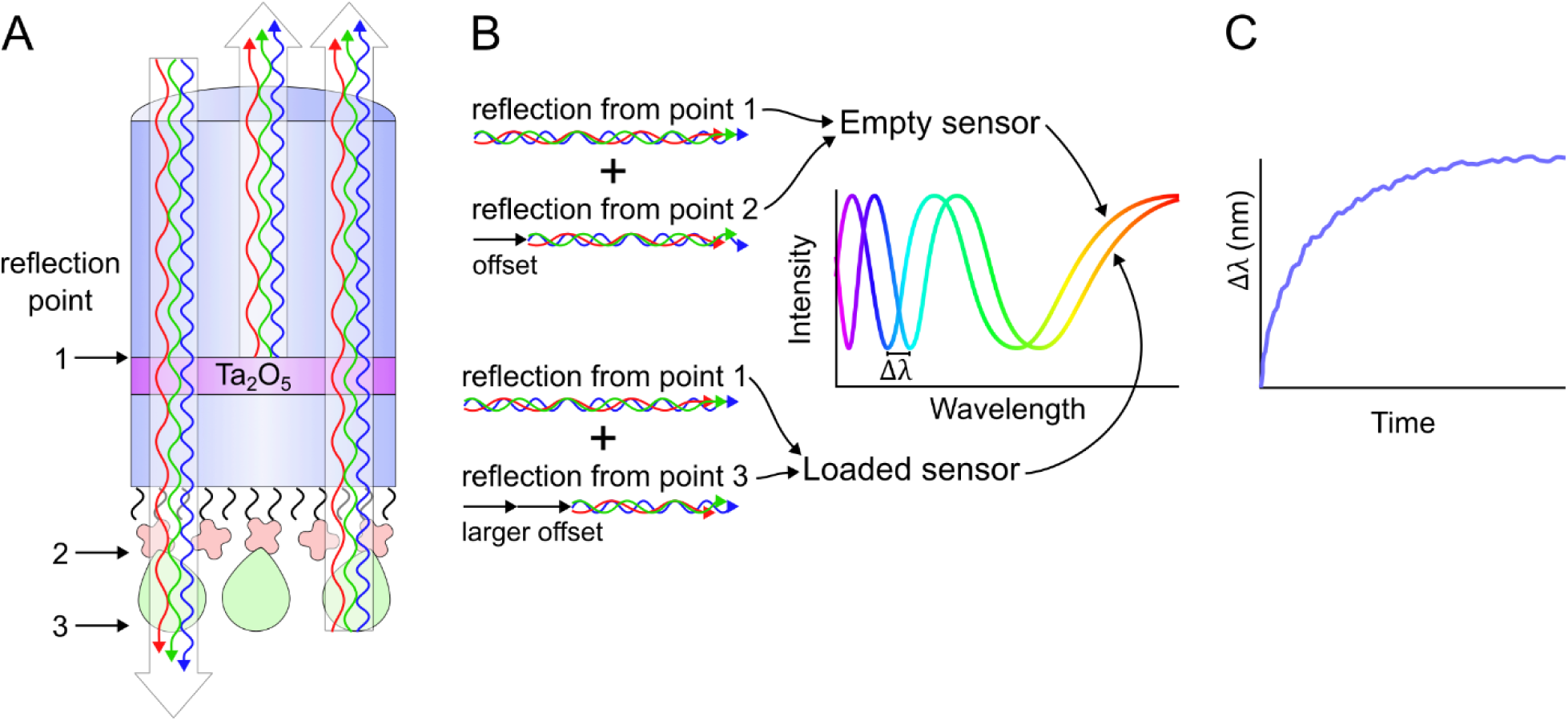
The physical basis of BLI measurements. A) BLI sensors are constructed from a fiberoptic piece (blue), a high refractive index film (often tantalum oxide, purple), a second length of fiberoptic material (blue), and a chemical coupling layer (wavy black lines). To this final layer, biomolecules such as protein antigens (red crosses) are attached. During an experiment, the sensor can bind to a test molecule (green teardrops). White light shone down the sensor is reflected back at 2 distinct points at a given moment. Reflection point 1 is the tantalum oxide film, and is built into the sensor. Reflection point 2 is the edge of the antigen and is changed to reflection point 3 when test protein is present. B) The different wavelengths contained in the white light undergo constructive or destructive interference based on the distance between the reflection points, creating an intensity pattern which shifts in wavelength upon binding. C) The wavelength shift can be plotted over time as binding occurs to measure the rate of biomolecular interactions.

The data provided by BLI is similar to that generated by the established technique surface plasmon resonance (SPR), which was first described in 1983 ^23^. Both rely on the reflection of light from a biomolecule-coated surface, but the physical basis for their measurements are quite different. Where BLI measures the phase difference in the interference pattern of reflected light, SPR relies on changes in total internal reflection of obliquely applied, polarized light to the back side of a metallic film loaded with biomolecules. Regardless of the mechanism, these two approaches provide nearly identical binding data that can be analyzed in much the same manner. In fact, BLI data can be imported into SPR software and analyzed without any modification whatsoever, and vice versa. This is a boon for BLI because it can fall back on the wealth of knowledge and experience established specifically for SPR experimental design and analysis ^24,25^.

### Definitions

The term **biosensor** is used to describe the device housing the surface to which a sample biomolecule is bound. For BLI, this is a small disposable biosensor tip (often shortened to “**tip**”), which is typically delivered in trays of 96 tips and generally cost between 5-20 dollars for each sensor, depending on surface chemistry, as of 2023. These tips replace the biosensor chips used for SPR, however SPR chips are generally much more expensive (several hundred dollars each) and include the gold-coated biosensor surface as well as microfluidic channels for directing fluid flow. Fiber optic SPR (FO-SPR) systems are now also available which forgo chips in favor of gold-coated tips, which are handled similarly to BLI tips.

By convention for BLI/SPR, the molecule bound to the surface of the biosensor is referred to as the **ligand**. This is notably different from standard biochemical terminology, where a ligand is a small soluble molecule which binds to a typically larger receptor. The ligand may be any type of molecule (protein, nucleic acid, lipid, small molecule, etc.), but is most often a protein such as an antibody or antigen, and it is always attached to the surface of the biosensor prior to binding measurements. Other proteins, such as streptavidin, may be used as capture reagents for the ligand, but generally only the molecule of interest is considered a ligand whereas the streptavidin in this example would simply be considered part of the biosensor. The soluble component which is varied in concentration is referred to as the **analyte**, and this may also be a biomolecule of any type. Selection of which molecule will be the ligand, and which will be the analyte will be discussed further in a later section, but depends on several factors including the size, number of binding sites, and stability over a wide range of protein concentrations/conditions.

Response refers to the magnitude of signal observed at any given moment by the BLI sensor, and a plot of this data over time is called a **sensorgram** (sometimes sensogram). The units of response in BLI are nm and refer to the wavelength shift described in figure 1. Critically, the response value is linearly proportional to the mass attachment to the tip, with some limitations such as buffer composition and analyte type. For practical purposes, a protein-protein interaction will show a linear relationship so long as the buffer composition (salinity, pH, solvent/solute admixture) is held constant.

Binding **affinity** describes the strength of a bimolecular interaction. This can be quantified in several different ways including a 50% effective concentration (**EC_50_**) from an ELISA or a dissociation constant (***K*_D_**) from a BLI experiment. EC_50_ values are generally easier to determine because the observed effect can be from any assay that produces a dose-response curve such as an ELISA or cellular assay, but EC_50_ values cannot generally be compared between different assays. *K*_D_ measurements, however, directly assess the binding strength of a complex, and are (ideally) completely assay independent. The technical definition of *K*_D_ is as follows: given molecules A and B form a complex AB (reversible reaction shown in **Equation 1**), and provided a large excess of molecule A, what is the concentration of B at which half is unbound (A+B) and half is complexed (AB).

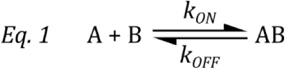

Bimolecular binding technically follows second-order kinetics, but by using a large excess of molecule A, we can hold that value constant over time. This allows us to assume pseudo-first-order kinetics, which has a much simpler mathematical description. *K*_D_ describes the equilibrium state of a binding pair and can be used to calculate the fraction bound for a given concentration of each component, but it can also be thought of as the point at which the rate of binding is matched by the rate of dissociation. These relationships are shown in **equations 2**.

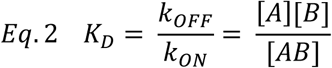

The equilibrium dissociation constant *K*_D_ has units of molarity (M) and can be measured using end-point assays, but it is often useful to know the rate constants, which can only be measured using real-time kinetic assays such as BLI. The dissociation rate constant (***k*_OFF_**) has units of per second (s^-1^) is frequently also written as (***k*_d_**), which is distinct from the equilibrium dissociation constant *K*_D_. Likewise, the association rate constant (***k*_ON_**) has units of per molarity second (M^-1^s^-1^) and is frequently referred to as (***k*_a_**), which is distinct from the equilibrium association constant (***K*_A_**). *K*_A_ is equal to 1/*K*_D_. Curve fitting of the BLI data allows direct measurement of *k*_ON_ and *k*_OFF_, which are then used to calculate *K*_D_. Curve fitting and parameter calculation will be discussed further in the analysis section.

### Experimental design

Measurement of binding kinetics by BLI requires careful experimental design, and there are several possible setups depending on the properties of the antigen-nanobody pair. This can also be directly extended to nearly any other binding pair of biomolecules, but for simplicity we will focus on the interaction of nanobodies with antigens here. The first important choice is the selection of using the nanobody or the antigen as the ligand versus as the analyte. This is a step of critical importance and there are several points to consider when making this selection including size, stability, and availability. BLI is often touted as a label-free analysis method, but this is only partly true. Because one of the components must be attached to the biosensor surface, some kind of chemical modification of one of the components is often required. However, in practice, this is rarely a problem and the advantages of BLI tend to far outweigh the limitations for binding kinetics measurement.

Nanobodies are relatively simple to use as either ligand or analyte in BLI because they are generally stable and consistently sized, they have a single binding site, and they are easy to label. Traditional antibodies and multivalent nanobody constructs should be used preferentially as ligands because bivalent analytes exhibit increased binding due to avidity effects and require more complicated analysis. This is discussed further in the analysis section.

Antigens are typically more varied in terms of their size, properties, and stability. Because of this, the antigen will generally determine which assay designs are possible. When possible, the antigen should be the ligand. The primary reason for this is material use, as most experiments use 100-500ng of ligand per tip, whereas analyte usage varies dramatically depending on the affinity and size anywhere from about 10pg per sensor (for a 15 kDa protein with a 1 pM *K*_D_) to over 300 µg per tip (for a 500 kDa protein with a 1µM *K*_D_). When testing a large number of proteins (such as a list of nanobody candidates from a screening experiment) against the same antigen, it is also beneficial to have a single batch of antigen of well characterized quality, concentration, and labeling efficiency in order to ease comparison between different nanobody clones. Conversely, some antigens do not make suitable ligands for a variety of reasons including low stability, oligomerization, difficulty labeling, or just poor performance in pilot experiments. For this reason, it is often useful to attempt both orientations.

#### Biosensor selection

Biosensors are produced commercially by device manufacturers. This manuscript will focus on the OCTET system (presently owned by Sartorius) as it is the most readily available system for most users at the time of writing, but most of the recommendations made here will be generally applicable to any BLI device. The currently available biosensors fall under a few major categories: antibody capture, affinity tag, and chemically defined. While successful nanobody-antigen interaction studies can be carried out using a wide variety of biosensors, we have found that streptavidin sensors were the most useful for the widest variety of possible experiments.

##### Antibody capture

Antibody capture tips include a wide array of isotype-specific Fc capture reagents as well as protein A, G, etc. These sensors are set up specifically for antibody ligands, but almost always target the Fc regions which are not present on nanobodies. It would be possible to construct a makeshift nanobody capture biosensor by combining a different biosensor with a high-quality commercial anti-VHH antibody such as Jackson Laboratories’ anti-alpaca VHH domain (128-007-232), but this would not be the generally preferred method. Another drawback of these sensors is the finite affinity of the capture antibodies for the ligand, which will lead to measurable dissociation over the course of an experiment. However, the fragility of the capture antibody-ligand interaction can also be exploited to refresh the sensor at the beginning of each cycle by regenerating and re-loading with fresh ligand. Further, affinity capture tips such as these may be used to capture ligand from a complex mixture, potentially allowing the use of unpurified sources of ligand such as whole blood, culture media, or lysates. Because BLI does not contain microfluidic components, there is no concern of clogging, though surface fouling and non-specific binding should still be considered when using unpurified biosamples.

##### Affinity tag

Affinity tag tips utilize multiple chemistries to target common protein tags such as GST, His, and biotin. These include capture antibodies such as anti-GST, which will behave similarly to Fc capture tips but will have specificity for appropriately tagged recombinant proteins. His-tagged proteins can be targeted with tips either containing a anti-HIS antibody that binds specifically to his tags, or through Nitrilotriacetic acid (NTA) tips which will bind to His tags similarly to Ni-NTA resins commonly used for recombinant protein purifications. We have found that the anti-HIS antibody has much better performance with his-tagged proteins, mainly through greatly decreased background protein binding compared to NiNTA tips. GST and his tag targeting tips may be washed and regenerated similarly to antibody capture tips, with the same benefits and drawbacks. If the GST tag approach is used it is critical that biophysical properties of the tag of interest is taken into account, this is especially critical if there is any potential for oligomerization, as GST tags can drive non-native dimerization.

Also in this category are the streptavidin tips, which are somewhat unique because of the exceptionally high affinity of streptavidin for biotin, about 10^-15^ M ^25^. This interaction is highly stable to changes in pH, addition of chaotropes, and extensive washing. This makes the connection behave, for practical purposes, similar to a covalent linkage. One can therefore wash and regenerate the sensor between cycles without needing to re-load with fresh ligand. The permanence of the streptavidin-biotin interaction also reduces baseline drift due to ligand desorption to negligible levels.

The primary challenge, then, is the addition of biotin to the ligand. This can be easily accomplished with commercial kits using amine-coupling or cystine-labeling, or genetically with an avitag. If using amine-coupling, it is beneficial to use a chromophore-containing reagent, for example Vector Laboratories’ ChromaLINK biotinylation kit. This allows quantification of biotin incorporation, which helps minimize batch variation and targeting a specific molar labeling ratio. This method modifies surface lysines as well as the N-terminus, the number of which will vary depending on the ligand. The ideal amount of labeling is one biotin per protein, but as amine-coupling is a random process it is important to acknowledge that the labeling will follow a poisson distribution. This contrasts with avitag labeling which results in site-directed labeling at the location of the tag with exactly one biotin per protein, depending on the efficiency of the biotinylation reaction. Site directed labeling may seem then, to be the preferable option, however one must also consider that labeling of any kind, including at the C- or N-termini can result in alteration of protein structure and function, possibly affecting binding, so one must take into account the pluses and minuses of site directed versus stochastic labeling for any given ligand. Site directed labeling will result in a single population of modified ligands whereas random labeling will give a mixed population with labeling at different locations, unless a large excess of reagent is used to drive the labeling reaction to completion. The site of labeling may even affect the orientation of the ligand on the biosensor surface. Because BLI is a surface adsorption-based method, it is possible for ligands to exhibit bias in which surfaces interact with the surface, and which surfaces are exposed for binding. Therefore, it may be beneficial to perform multiple different antigen labeling methods during pilot experiments to empirically determine the ideal method for each nanobody-antigen pair.

##### Chemically defined

Chemically defined tips include aminopropylsilane and amine-reactive tips. These tips do not contain capture proteins, and simply have a base layer of chemical moieties. This allows the most flexibility in labeling at the expense of needing to perform substantially more loading optimization. For this reason, chemically defined tips are not recommended for beginning users and should only be used when other options are not available for a particular ligand. For instance, amine-reactive tips are coated with carboxylic acids which can be coupled to a protein using EDC/NHS chemical methods. This is similar to the amine-coupling biotinylation reaction described above, and it is generally easier to perform one large biotinylation reaction and freeze small aliquots of protein with well-defined labeling efficiency than it is to perform a fresh amine-coupling reaction at the start of every BLI experiment. Additionally, it is critical to confirm ligand purity and quality prior to this kind of labeling because general chemical coupling will equally attach the ligand of interest as well as any proteinaceous contaminants or other primary amine-containing molecules.

#### Step types

After selection of biosensor type and ligand-analyte orientation, the next step is to design the steps of the experiment. There are several different step types which can be defined in the OCTET Data Acquisition software, and these should be broadly applicable to any BLI. An example sensorgram is shown in **Figure 2**, which represents a simplified experimental design incorporating each of the step types: biosensor preparation, blocking, activation, loading, quenching, baseline, association, dissociation, and regeneration.

**Figure 2.**
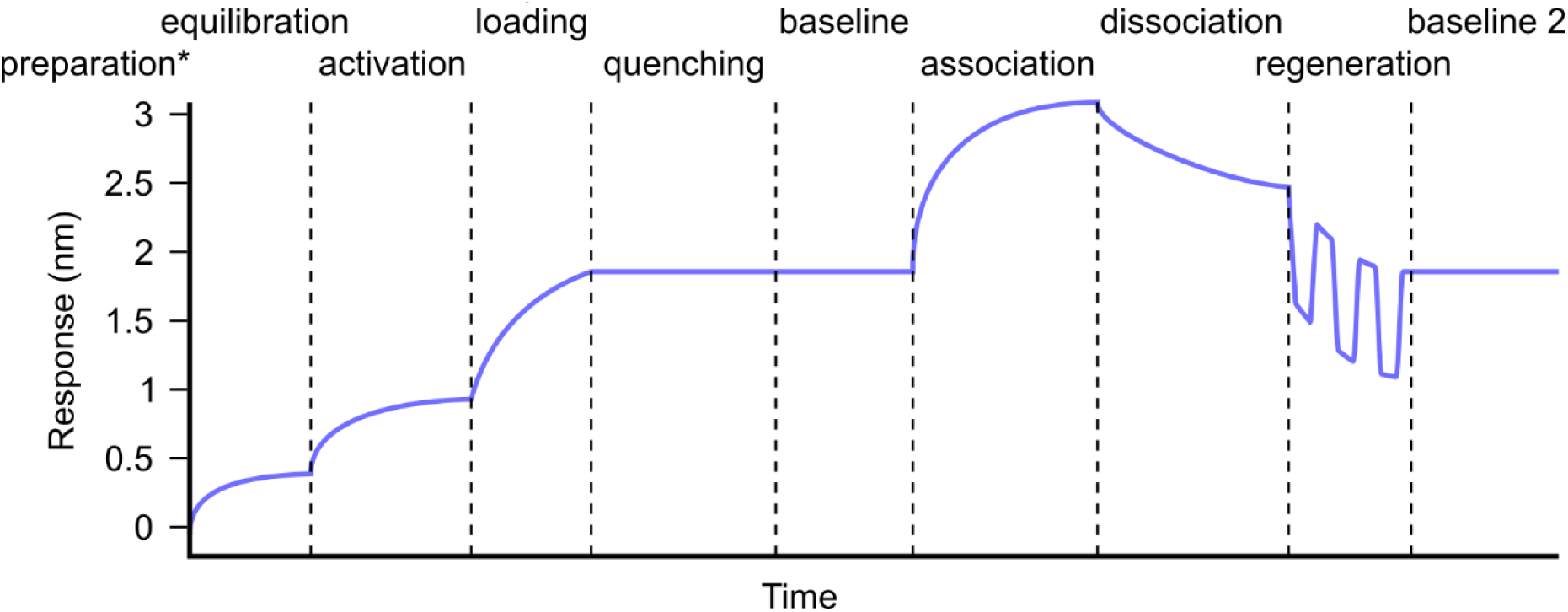
Prototypical sensorgram. Each of the basic step types including antigen preparation (*occurs prior to start of the experiment), blocking with running buffer, chemical activation (only some sensor types), antigen loading, quenching (only some sensor types). These steps are generally performed once during experiment setup, then followed by kinetic measurement cycles composed of baseline, association, and dissociation steps with a regeneration cycle in between each measurement cycle. Response values depict average expected magnitudes for each step type rather than binding of a specific molecule.

##### General considerations

Some settings apply to the entire experiment and can have a substantial effect on the outcome such as temperature, shaking speed, averaging, sensor offset, and total assay time, and well volumes. Temperature is critical parameter because *K*_D_ is temperature dependent due to its relationship with Gibbs free energy ^26,27^. For proteins, this relationship is highly complex and non-linear, and is modified by the nature of the non-covalent interactions that drive binding. This will mainly be controlled by the relative entropic and enthalpic contributions to the free energy of complex formation. Hydrophobic interactions tend to decrease at low temperature while charged interactions and hydrogen bonds are less effected, and this leads to increased, decreased, or equivalent *K*_D_ over a temperature range depending on the dominant interaction for a particular antibodies; and in fact, measurement of antibody *K*_D_ at a range of temperatures can be used as a method of identifying high quality candidates ^28^. Most OCTET systems have a very narrow temperature range of ambient plus 4°C up to a total of 40°C, and the default setting is 30°C. This is standard for the field at the time of writing, but may change as device capabilities improve.

Plate shaking is an important feature of BLI which limits the effects of diffusion rate on binding results. We recommend a shake speed of 1000 rpm for every step. Reducing this can slow binding, which can be used for loading high concentration samples, but it is preferable to dilute samples than attempt to slow loading by limiting the diffusion rate.

Averaging is a description of the number of read cycles which the OCTET averages to generate the data points. The inherent measurement frequency varies between equipment, and the number of cycles which is averaged ends up having relatively little impact on the final results in practice because it does not change the actual data acquisition rate, just the amount of averaging that is done in the raw data.

Likewise, the offset parameter does not typically need to be adjusted unless an unusually sized multi-well plate is used. Smaller offsets move the tip closer to the bottom of the well. 4mm is generally recommended, but 3mm can be used for tilted bottom plates. Greater sample volume is needed to adjust for increased offsets, which may slightly reduce the intensity of reflected light from the bottom of a flat plate.

Total assay time is an important consideration, primarily due to evaporation. *k*_ON_ is linearly proportional to analyte concentration, so excess concentration due to evaporation will tend to increase the apparent *k*_ON_, and decrease the apparent *K*_D_ by the same factor. In our experience, assays at 30°C should be kept to under 2 hours whenever possible, however increasing the sample volume may allow longer experiments. Little objective data exists for the evaporation, as this is dependent on temperature and humidity, but the authors have observed evaporation sufficient to disrupt data collection after as little as 4 hours using a 384 tilted well plate loaded with 40µL, running at 30°C with a 4mm offset. This includes the full time from pipetting samples into the plate until the final time a biosensor is dipped into a sample. Another critical feature is determining the sample stability (both of the ligand and the analyte), as assay results will be strongly disrupted if there are any issues with sample fouling over the total assay time. Replicate experiments should be conducted at the beginning and end of any assay window to make sure this is not an issue.

The well volume used for an experiment is dependent primarily on the plate type used. It is impossible to know what the minimum volume is for an arbitrary plate without testing, but generally half of the maximum capacity is a good starting point, however changing sensor offset may alter this slightly. Black plates are recommended to prevent reflections from the well bottom, and a glossy finish on plates is to be avoided. 96 and 384 well plates both work well, but the minimum possible sample volume of 40µL can currently only be achieved using OCTET tilted well 384 plates. Good alternatives plates are the Greiner Bio-One black flat-bottom polypropylene plates (96 or 384 well) 781209 (384), 655209 (96).

##### Biosensor preparation

OCTET biosensors require extremely careful handling. They should be maintained in their sealed original packaging for as long as possible, primarily to avoid moisture, dust, and unnecessary contact with the biosensor surface. At the time of writing, biosensors are shipped with a proprietary coating (very likely plain sucrose) which must be fully dissolved prior to beginning any experiment. This is accomplished by filling the biosensor soaking plate with 150-250µL of deionized (DI) water or buffer. The authors recommend DI water at this step for streptavidin tips, but a neutral isotonic saline like PBS may be more appropriate for sensors pre-loaded with antibodies or other delicate proteins. Assay running buffer may also be used here, but it is generally unnecessary.

Biosensors can then be loaded carefully into the overlaid sensor tray such that they contact the liquid. Ensure that the tip end of the biosensor does not come into contact with any solid material, including the side of the sensor rack. Tips should be soaked for at least 10 minutes, preferably 20 minutes, prior to starting a run. Only tips being used in the current experiment should be soaked, and they should generally be used the same day that they are prepared.

Tips can be saved for later use by dipping briefly into a 15% sucrose solution, dried at ambient temperature, and returned to a sealed package with desiccant. This technique can be used to pre-load a batch of tips with a particular ligand and stored for later use. However, it is generally preferable to load each experiment separately using a fresh aliquot from a single batch of labeled ligand.

##### Equilibration

The most common first step of a BLI experiment is to block the tip briefly in the assay running buffer. This step serves two purposes. First, it is useful to observe the behavior of each tip as it is introduced from plain water into the running buffer. The equipment will auto-zero each trace prior to collecting any data, and any defective tips will generally be clearly visible at this step. In our experience, defective tips are very rare, but improper preparation or mishandling may increase the chance of observing unusual results. In our standard running buffer, we generally see 0.4-0.5 nm of signal after 30 seconds. Second, it is useful to pre-bind the sensor with blocking compounds to prevent non-specific binding at the loading step. This step should be delayed until after loading if using a chemically defined tip.

This step is where running buffer is first introduced to the tips, and all tips of an experiment should have the same running buffer. This will require tuning for specific needs, but a good general starting buffer is 10mM HEPES, pH 7.5, 150mM NaCl, 3mM EDTA, 0.005% Tween-20, and 0.1% BSA. Changes to the buffer recipe should be brough forward into every step that uses running buffer, because relatively small changes can shift the observed signal significantly. Non-neutral pH (up or down) typically shifts the signal lower, while increased salinity typically increases signal. This is likely due to changes in the packing of the silane molecules used to coat the raw biosensor surface. However, because of referencing, it is not usually problematic to change buffer conditions to whatever is required for a particular binding interaction. Streptavidin sensors can handle harsh buffers and wash conditions, but it should be noted that many other tip chemistries will dissociate their ligands, and possibly damage their capture molecules if exposed to overly harsh conditions.

##### Activation

Activation only applies to amine-coupling tips and involves incubation with a mixture of EDC and sulfo-NHS. This can be acquired as a kit or separately with similar effect, but amine-coupling should not be one’s first choice for tip chemistry. One should expect approximately 0.3-0.5 nm of signal at this step, but it will depend on concentration and the exact compounds used for activation.

##### Loading

Loading is one of the most critical steps of each assay and should be optimized with pilot experiments. Greater loading densities will linearly increase the amount of signal achieved during association steps, but overloading can lead to crowding or mass transport limitation. Proper loading should be approached empirically, but 1-2 nm of binding is often a good starting point. Actual capacity of biosensors will vary substantially based on the tip chemistry, and small variations will also be seen between individual tips, even from the same package. This will result in different rates of loading as well as signal at saturation. A good starting point is 10µg/mL for loading. Results will vary depending on tip chemistry and batch, but an example experiment with streptavidin 16 tips loaded with identical ligand solutions of a biotinylated (∼2.5 biotin/protein) 26 kDa protein over 265 seconds reached an average signal of 2.84±0.42 nm (%CV = 5%). This is approximately in line with the manufacturer’s certificate of analysis. For streptavidin tips, ligands with excessive biotins per protein will reduce tip capacity, as will proteins contaminated with free biotin.

The signal obtained from a defined amount of mass attached to a biosensor should be theoretically calculable for an arbitrary ligand-analyte pair based on their relative molecular weights, the stoichiometry of binding, and the amount of signal seen during loading. For example, if a 17 kDa nanobody is loaded to 1 nm then one would generally expect to require approximately 5 nm of loading for a 150 kDa bivalent antibody in order to achieve the same level of signal from the same analyte.

For reference, real world examples from experiments we have conducted with streptavidin tips gave roughly the following signals: A biotinylated 26 kDa protein with a single binding site was loaded to 3 nm, a 17 kDa nanobody reached 0.7 nm of binding. A neighboring tip showed a lower capacity and only achieved 2.25 nm of binding, and as a result only reached 0.5 nm of binding with the same concentration of nanobody. Next, an 87 kDa bivalent nanobody construct was tested with the same batch of ligand. In this case 1.8 nm of ligand loading led to 1.1 nm of bivalent nanobody binding. Next, using a 700 kDa trimeric ligand loaded to 0.75 nm led to 0.02 nm of monovalent nanobody binding. These can be compared by correcting the response signal by the molecular weight for each the ligand and the analyte. We can then get a unitless ratio of analyte signal (nm/kDa) per ligand signal (nm/kDa) and we get values between 0.33 and 0.36 after correcting for the stoichiometry of each interaction. The amount of analyte used in each case was over 100 times the *K*_D_. This ratio appears to be relatively stable over a handful of nanobody-antigen pairs with different stoichiometry in both orientations, but we have not tested it exhaustively and it should be used only as a guide for planning pilot studies, or to help estimate whether a particular ligand-analyte pair is likely to give good results. Similar ratios ranging from 0.33 to 0.45 can be calculated from BLI traces published by other studies where both full sensorgrams and approximate molecular weights are available ^29–33^. A physical interpretation of this ratio may be that approximately one third of loaded ligands are able to bind an analyte, but further testing would be required to prove this. It would be reasonable to expect that this ratio will change based on the quality and purity of the loaded ligand as well as the final loading density.

##### Quenching

Quenching is necessary to fill any remaining binding sites not used during loading. If left active, these extra sites can cause analytes or other compounds present in samples to bind during later steps, confounding the results. This can apply to either chemical or biochemical loading methods. For chemical reactions such as amine-coupling, quenching should chemically inactivate any remaining succinimide esters by reacting with any primary amine-containing small molecule including ethanolamine, tris, or glycine.

For streptavidin tips, loading is rarely complete, and a substantial number of free binding sites will remain. This can be mitigated by filling all remaining sites before proceeding to kinetic measurement cycles. For streptavidin, dilute solutions of free biotin are sufficient to rapidly and permanently saturate the surface. This is only necessary when biotinylated proteins will be present in later samples, but many biosamples do contain trace amounts of biotin and biotinylated proteins. This quenching will persist through regeneration.

This type of quenching/blocking should be approached with caution for other tip chemistries, but may be helpful in some cases. Most other tip types bind to a protein rather than a small molecule such as biotin, and therefore any compound used for this purpose will be similarly bulky as the ligand of interest. Thus, blocking the remaining binding sites could result in overly dense loading of the sensor surface and steric hinderance of the ligand, which must be strictly avoided. In these cases, it would be preferable to ensure that no interfering proteins or compounds are present in the buffer or analyte solutions.

##### Baseline

The baseline step is beneficial for confirming the level of drift as well as providing a starting point from which to calculate the association and dissociation. A 30 second baseline is generally sufficient. Consistent drift can be removed by reference subtraction, but large amounts of drift will negatively impact the results regardless of proper reference subtraction if it is due to ligand dissociation as the response value at saturation (**R_max_**) value will vary over the course of a single cycle. This can be a key step to optimize the type of attachment to the sensor, as if a given approach has significant drift (mainly due to ligand loss from the sensor) this is a good reason to switch approaches. Empirically we have found that the streptavidin tips perform much better compared to anti-GST or anti-HIS approaches in this regard.

##### Association

The association step is used to calculate *k*_ON_ and is dependent on accurate knowledge of the analyte concentration, and any underestimation at this stage will lead to falsely low (strong) *K*_D_ values. The appropriate concentration range for kinetics measurement is 0.1–10×*K*_D_ with at least 5 concentrations over that range. In order to prevent signal spikes due to changes in buffer composition, the stock protein should be sufficiently diluted in the running buffer. For buffers of similar composition, 100-fold is generally a sufficient dilution factor to avoid buffer spikes. If spike still occur, then dialysis can be used to equalize conditions somewhat, but additives like Tween-20 and BSA are not generally possible to dialyze, and low CMC surfactants and low molecular weight blocking compounds may be substituted, but should be tested. A better strategy is typically to further purify the analyte, exchange into running buffer base without the large molecular weight additives, then dilute at least 100-fold as normal.

As the *K*_D_ is often not known prior to BLI, an ELISA EC_50_ is a good starting approximation, and a wider 10- to 100-fold dilution series can be used to home in on the proper range. By definition, at the *K*_D_ the curve will reach equilibrium at half of R_max_, if given enough time. Equilibrium need not be reached, however unless performing equilibrium analysis specifically, which is only ideal for very fast or low affinity analytes. Minimum step times are dependent on *k*_ON_, but there must be curvature present in order to make fitting possible. Ideally, the highest concentration tested should be allowed to level out. For nanobodies, typical *k*_ON_ values range from 10^5^ to 10^6^ M^−1^s^−1^, but somewhat counterintuitively the primary factor affecting equilibration time is the dissociation rate *k*_OFF_, which typically ranges from 10^−2^ to 10^−4^ ^34,35^. Given the slow off rates of many nanobodies, it is often impractical to wait for equilibrium to be reached, however it can be calculated using **Equation 3** ^24,36^.

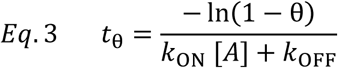

Where θ = 0 represents starting conditions and θ = 1 represents steady state and [A] is the concentration of analyte. This is concentration dependent, and higher concentrations will reach equilibrium faster. Assuming we wish to reach 90% of equilibrium binding for the middle concentration where [A] = *K*_D_, it will take just under 2 minutes for a nanobody with *k*_OFF_ = 10^-2^ s^-1^, just under 20 minutes when *k*_OFF_ = 10^-3^ s^-1^, and over 3 hours when *k*_OFF_ = 10^-4^ s^-1^. Because 10^-4^ s^-1^ is a relatively common *k*_OFF_ for nanobodies, it is important to consider this factor. Even if we look at the 10×*K*_D_ sample, it will take 600 seconds to get halfway to equilibrium. In such cases, it may be prudent to include an even higher concentration sample, because a 100×*K*_D_ sample will reach 90% equilibrium in just under 4 minutes. Because a nanobody with *k*_OFF_ of 10^-4^ s^-1^ is likely to have a *K*_D_ between 100pM and 1 nM, it is safe to increase the sample concentration up to 100×*K*_D_ = 10-100 nM without needing to worry too much about non-specific binding.

Non-specific binding becomes much more likely as analyte concentration increases. In general, small proteins like nanobodies (12-17 kDa) can be used up to 100-300 nM without substantial effects, but this may not be true for larger macromolecules. In our experience, concentrations around 1µM are where non-specific binding become common. If non-specific binding is observed in pilot studies or otherwise anticipated, the running buffer can be modified, concentrations decreased, or offending substances removed. For severe non-specific binding which might arise when studying low affinity interactions, additives such as 0.6M sucrose may prove useful ^37^.

Two association steps can be performed back-to-back for competition experiments. This is useful for general competition experiments as well as epitope binning. In either case, it is important to consider whether to include the analyte from step 1 in the buffer for step 2. In the case of high affinity nanobodies it is likely not necessary to include analyte 1 in the second buffer, but for faster off rates it may be obvious that the first analyte is dissociating during the second binding step, complicating the results. If included analyte 1 should be kept at the same concentration as in step 1, to avoid shifting the equilibrium. Further, the shape of the curve for step 2 should be observed carefully. For slow off-rate nanobodies, a rapid increase in signal should always be due to binding at a non-overlapping site, but for fast off-rate nanobodies/proteins one should consider that analyte 2 is exchanging with analyte 1, but still binding to site 1. Whether this is occurring can be reasoned based on the direction and amount of signal shift compared to the relative molecular weights of the two analytes. Allosteric binding by analyte 2 followed by conformational changes that prevent binding at site 1 is also possible. Prior to performing this type of experiment, one should ensure that analyte 1 is fully saturating and at equilibrium. 100×*K*_D_ is usually sufficient for this, and should be continued until the curve flattens.

##### Dissociation

The dissociation step is always performed immediately following an association step, and it is used to calculate the dissociation rate constant *k*_OFF_. This step is generally best performed in the same well that the baseline step was performed in. When possible, a fresh running buffer well should be used for each cycle. Baseline, association, and dissociation should be performed in the same plate with the same well volume, to prevent changes in reflection/refraction from introducing additional noise. If throughput is a concern, then the same buffer well can be re-used for multiple cycles, with the only concern being contamination from previous cycles. However, the amount of carryover is generally minimal, and we have not observed noticeable changes with repeated use. As a precaution, we suggest starting with the lowest concentration and increasing with each cycle. The manufacturer claims that a fully saturated tip will transfer 10^9^ molecules, resulting in an 8.3pM solution assuming 100% dissociation and a 200µL well volume. This ignores liquid carryover on the tip, but also likely assumes 100% saturation during the loading step as well. All together, our observations suggest that this is likely accurate, but the primary reason to re-use wells is to save space in crowded layouts and we would generally consider using fresh wells to be best practice.

Dissociation should generally be allowed to reach at least 5% signal reduction in order to get an accurate reading. While the absolute rate of decrease will occur faster for higher concentrations, all concentrations will decrease proportionally, so increasing the concentration during association will not result in reaching 5% reduction faster. The time required at this step can be calculated using **Equation 4**, derived from the equation for one-phase decay, where θ is the fractional progress towards steady state ^36^.

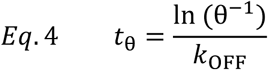

Using 0.95 for θ gives the time to dissociate 5% from R_max_. For a *k*_OFF_ of 10^-4^, it will take 8.5 minutes to dissociate 5%, 1.4 hours for 10^-5^, and 14 hours for 10^-6^. It is clearly impossible then to definitively show an off rate of 10^-6^ given the time limitations imposed by evaporation, but this is not a hard cutoff and depends on how clean the curves are. The lowest *K*_D_ that the OCTET software will assign is 10^-12^ (1pM), which is a *k*_ON_ of 10^6^ M^-1^s^-1^ and a *k*_OFF_ of 10^-6^ s^-1^, below this it will report >10^-12^ M. The time required for this step is unfortunately a physical limitation of dissociation kinetics and cannot be easily circumvented.

One strategy for reducing the total experiment time, is to perform short dissociation steps for the lower concentrations and only perform a full-length dissociation step for the highest concentration tested. This works because the dissociation rate is not dependent on concentration or saturation level of the biosensor. The highest concentration is used because the higher starting point yields a larger absolute change in response over time, and therefore improved signal-to-noise of the resulting drop. However, some software, such as OCTET Data Analysis HT 10, struggles to analyzing data with variable step times for each cycle.

##### Regeneration

Regeneration is typically a series of steps designed to remove excess analyte from the last completed kinetics cycle and re-set for a new cycle. This is simple for some tips and ligands, but nearly impossible for others. The general strategy is to find conditions which cause the analyte to dissociate rapidly without irreversibly damaging the ligand. The most common regeneration solution is low pH glycine, between pH 1.5 and 2.5, but this also needs to be tested empirically. These conditions work well for most, but not all, nanobodies. We recommend three cycles of 15 second dips each into 10mM glycine pH 1.7, then into running buffer. The reason for repeated short cycles is putatively to limit irreversible unfolding of the ligand, but this is often unnecessary and single stage regeneration is also generally acceptable.

The extent of regeneration is verified by checking the level of baseline after regeneration, which should match very closely the signal of the previous baseline step. Small shifts up or down are common, but not ideal. **Figure 3A** compares complete versus incomplete regeneration. Residual ligand-analyte complex will alter the curves as the pre-existing complex is free to dissociate similarly to the newly formed complex. This can be accounted during analysis, and is exploited in a process called kinetic titration (or multi-cycle kinetics), but currently available OCTET software is not capable of this type of analysis. This will be discussed further in the analysis section.

**Figure 3.**
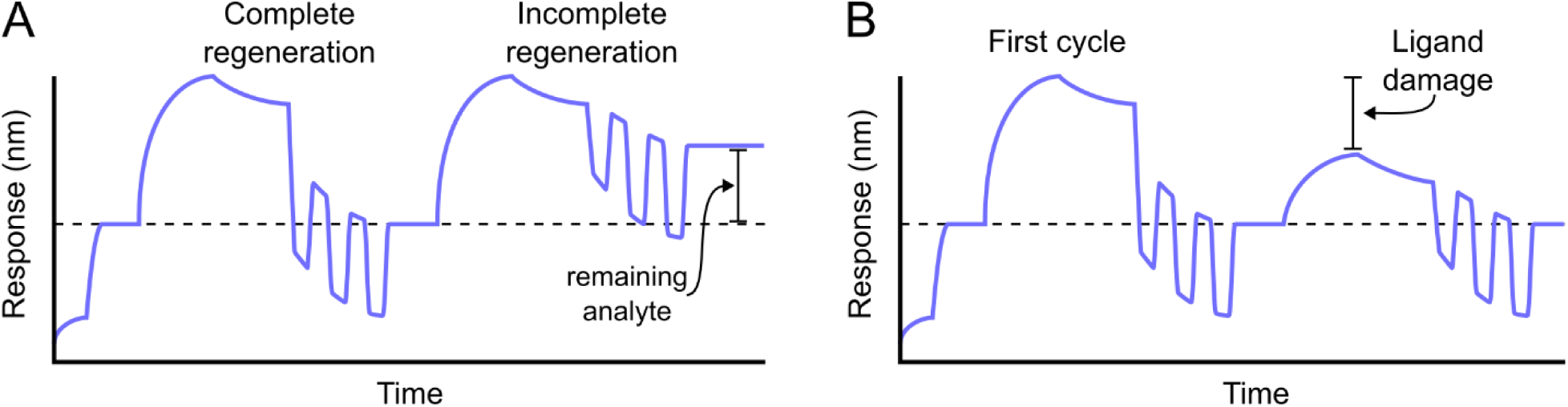
Evaluating problems with regeneration. A) Sensorgram of a simple experiment with a initial blocking step and ligand loading followed by a measurement cycle (baseline, association, and dissociation). The response does not return to baseline following dissociation, so regeneration is required. The first cycle shows complete regeneration, which rises and falls rapidly depending on buffer composition, but returns to the initial baseline value upon completion. Following the second cycle, the response value remains above the initial baseline, indicating that some analyte remains bound to the sensor and regeneration was incomplete. B) A similar first cycle with apparently full regeneration is followed by a measurement cycle with the same analyte at the same concentration, but showing reduced response signal during the association step. This indicates that the ligand was partially damaged during the first regeneration step.

Conversely, some conditions may damage the ligand, reducing the capacity of the tip for analyte binding (**Figure 3B**). In theory, this does not alter the curve shapes, but care must be taken during analysis as each cycle will have a different R_max_, and it is possible that some ligand molecules will be partially damaged, confounding analysis even further by creating a heterogeneous surface. Using streptavidin tips, it is unlikely that ligand will dissociate from the biosensor surface, which would present as a decreased baseline response following regeneration. However, other tip chemistries may experience this, and Ni-NTA tips, for example, can be regenerated with imidazole solutions to completely remove the ligand from the sensor, which can then be re-loaded with fresh ligand before the next cycle.

In the case of incomplete regeneration or ligand damage, the time in or composition of regeneration buffer can be adjusted. Several different categories of regenerant exist which act in different ways, depending on the nature of the interaction (**Table 1**). Low pH is the most common condition and acts by protonating acidic residues and disrupting hydrogen bonds/electrostatic interactions while high pH deprotonates basic residues and also alters hydrogen bonding/electrostatic interactions. More exotic reagents can be used depending on the biophysical nature of the interaction, and a variety of different compounds can be tested beyond those Table 1. However, more extreme conditions risk damaging not only the ligand but also the biosensor surface, so should be attempted only after milder conditions have failed. Specific molecules should be used whenever possible, such as competitive peptides or small molecules, sugars for lectins, and imidazole for Ni-NTA. A tip may be used until its capacity has degraded, but the tolerance for degradation can be selected on a case-by-case basis depending on target difficulty and signal-to-noise achieved using the damaged ligand. Alternatively, kinetic titration can be used to avoid regeneration entirely.

**Table 1:**
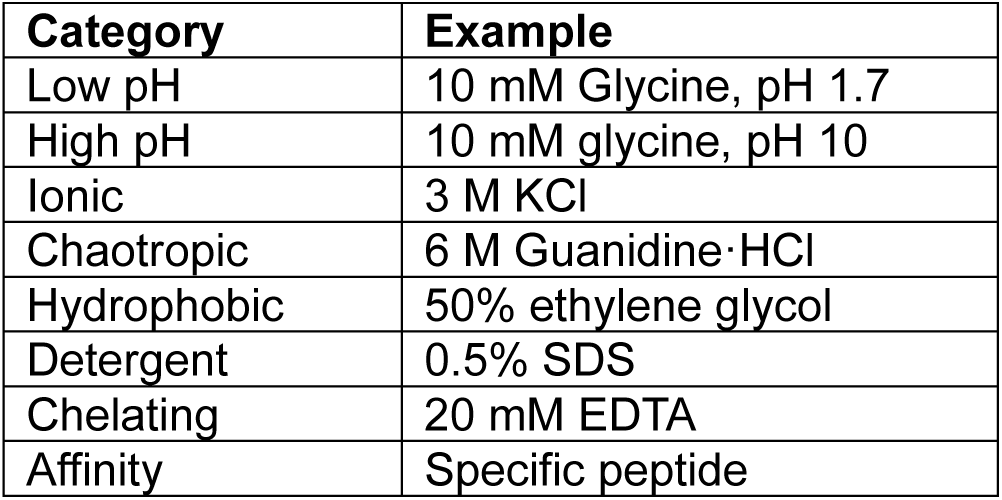
Example regeneration buffer ingredients.

### Controls

Artifacts in BLI data are unavoidable, but most common types can be effectively controlled for with minimal additional cost or effort. The most basic control is adding an addition sample with no analyte. This controls for differences in buffer composition between the running buffer in the baseline and association steps. Some buffer components such as DMSO have an outsized impact on signal, but ideally these would be tightly controlled in sample composition rather than relying on referencing to fix. This control can also control somewhat for drift, due to ligand dissociation from the sensor, though imperfectly, and drift is not an issue for streptavidin biosensors. The second, and more important, control is a reference sensor. This sensor gets loaded with either nothing or an irrelevant ligand, and it gets carried alongside the other sensor tips into identical analyte solutions. This helps control for non-specific binding as well as changes in refractive index due to the presence of large quantities of analyte. This control must absolutely be included in every experiment. Testing an analyte such as a nanobody or antibody against multiple different ligands can also help establish specificity. The preferred method, however, is called double-reference subtraction and involves running both of these references simultaneously. Because this requires no additional sensors, double-referencing is usually the right choice even though it provides minimal additional benefit over a reference sensor.

Replicates are also important to include when performing BLI. While pilot experiments may be performed with one sensor per condition, proper kinetic measurements should include at least two parallel sensors. Ideally, separate experiments should also be performed on separate days and with separate batches of protein. When using replicates, take care to keep the exact same experiment design including step lengths so that the analysis software can analyze the replicates together.

Additional controls may also be necessary if pilot experiments indicate complex binding phenomena. In general, if complex (not 1:1) binding is observed, then non-BLI/SPR method should be used to differentiate between the alternative explanations, but nevertheless there are some methods that can address specific cases. For example, by fully saturating the sensor for different lengths of time prior to dissociation, one can learn whether their interaction is affected by a ligand-induced conformational change ^38,39^. Strong mass transport limitation can be detected by observing a linear association phase which saturates without showing much curvature ^40,41^. Critically, however, the fitting of more complex models to existing data cannot reveal which model is correct. A more complex model will almost always fit the data better simply because it has more degrees of freedom, so further experiments are always needed to understand the molecular mechanims underlying complex binding.

### Experiment types

Multiple experiment types can be used to learn different pieces of information. This guide will cover four basic experiment types, which have substantial overlap in what they can teach us about a set of interacting biomolecules. Pilot experiments, full kinetics experiments, quantitative competition experiments, and epitope binning experiments. There are additional use cases for BLI which are beyond the scope of this guide, including quantitation experiments which use a variety of methods to determine the concentration of a particular analyte, however our focus is on experiments that are optimal for nanobody-antigen interactions. The kinetic and competition experiments used in this guide require access to high purity analytes and are used to biochemically characterize interactions.

#### Pilot

Pilot experiments are always the first step to developing a new BLI method. It is often not known what the approximate binding strength is prior to undertaking kinetics experiments, and values like the EC_50_ provide very limited guidance for setting experimental parameters. These experiments provide key information about whether an interaction is detectable using a given biosensor chemistry, loading conditions, or ligand/analyte identity and orientation. The most important information that needs to be obtained with a pilot experiment is the overall level of signal that is achievable for a given binding pair, the concentration of analyte that results in strong but not overwhelming binding, and approximate *k*_ON_, *k*_OFF_, and *K*_D_. Spending a few hours and a modest number of biosensors in a pilot experiment can save a significant amount of time and dozens of replicate biosensors from being wasted on full scale tests with suboptimal parameters.

#### Full kinetics

As stated earlier in the introduction, BLI is one of relatively few methods that can give binding kinetics for a biomolecule pair. Robust determination relies on testing multiple concentrations, and kinetics experiments should be designed to cover a broad concentration range. It bears repeating that only 1:1 interactions can be properly quantified using BLI, or any other currently available technique. However, more complex interactions can be monitored and compared relative to other similar interactions, and these experiments can be quite useful for understanding the biology of complex systems so long as it is correctly reported as non-quantitative.

One step under kinetics is steady-state analysis. BLI can use steady-state analysis to measure the *K*_D_, which represents the equilibrium dissociation constant. This should only be used where interactions are too rapid for proper kinetics experiments, and simply entails plotting the level of steady state binding on a dose-response curve, where 50% signal is equal to the *K*_D_. In general however, protein-protein interactions like nanobody/antibody-antigen interactions are too slow to reach equilibrium for this method to be practical. Fortunately, kinetic binding experiments provide more information than steady-state binding experiments.

#### Competition

Competition experiments can inform about whether two different analytes bind to the same physical location on a ligand. This is called the epitope when talking about nanobodies/antibodies, and finding multiple non-overlapping antibodies is of significant interest to many researchers. It can also be used to explore the relative binding affinities. Because BLI cannot distinguish the identity of molecules bound to the sensor, just the total mass bound, it is important to know the relative mass of the two competing analytes, and they can be bound consecutively in different steps, with this contribution to the signal able to be distinguished. The primary advantage of BLI competition experiments is the fact that data can be observed in real time. This added time dimension can help distinguish situations whether the second analyte binds because it is non-competitive or because the first analyte already dissociated. When properly set up, BLI is also actually faster than an equivalent ELISA experiment, because there is no need to wait for wash steps, reach an arbitrary end point, or develop signal. Like most other competitive binding assays however, BLI cannot distinguish between orthosteric and allosteric competition the same way that direct epitope mapping (HDX-MS, NMR, and crystallography/Cryo-EM) can.

#### Epitope binning

Epitope binning is a high-throughput method which is a natural extension of the competition experiments discussed above. Current protocols use an exhaustive round-robin approach to test every possible pairing from a set of analytes. This is most commonly used for screening applications and is named epitope binning because its primary use case at this time is to test a large pool of antibodies and place them into buckets based on whether they bind competitively. This saves time at later stages of biochemical characterization because one can simply obtain epitope information from one member of each bucket and the data will inform the approximate binding site of a whole class of antibodies. This is particularly useful when a set of commercially available antibodies with known epitopes can be compared to an unknown molecule in order to approximate its binding site. Epitope binning is not quantitative, but this is usually not important and any interactions that do need to be followed up on can be tested individually using more targeted approaches

### Data analysis

#### Software

BLI data can be easily analyzed using the bundled OCTET software, and this guide will be based on OCTET Data Analysis HT version 10.0. Newer versions are available (version 13 at the time of writing), but most previous versions are sufficient for basic kinetics analysis. No centralized database of software or feature lists exist, but basic kinetic analysis has been possible in every version to date, competition experiments can be performed and analyzed using every version to date (albeit with some manual post-processing), and epitope binning has been available as an officially supported feature since version 8.0 (released around 2016). HT stands for high-throughput and the HT version of the analysis software was released along with version 9.0, and while there is a standard version of the software, there is no notable benefit to its use in its current form. The Data Analysis HT software is not the only option however, and raw or partially processed data can be exported and analyzed in other software such as Graphpad Prism, SPR analysis software like BIAevaluation, or other custom software like TitratonAnalysis ^42^. The standard Octet software will be sufficient for most new users, but there are several key features it currently lacks, such as the ability to analyze single-cycle kinetic titration experiments. To summarize, the experiments discussed can be reasonably performed on any BLI equipment using whatever software version is already set up for use with that equipment. The precise methods of analysis may differ slightly, but the only key features necessary for the performance of these protocols are basic reference subtraction and curve fitting. More advanced experiments such as competition and epitope binning are easily interpreted manually, however current generation software features improve the ease and repeatability of these analysis.

It is fortunate that the quantification of binding rate has been made relatively simple by modern computing, and no longer requires log transformed Scatchard plots. Because simple 1:1 binding can be fit using pseudo-first order kinetics, we can use non-linear regression with a single exponential for each the association and dissociation steps. Further, this work happens in the background of most common software. Nevertheless, it is important to understand what calculations are being performed on the data, and there are several stages of data analysis that must take place to get reliable quantification. Critically, all of these steps are identical to the processing steps necessary for SPR analysis, which is the reason why these data are interchangeable during quantification.

#### Referencing

The raw sensorgram data does a good job of intuitively showing the amount of binding, but the units are relative and need to be converted to a form that can be more easily fit to a model. This starts with background subtraction. Using double reference subtraction, we first subtract the primary baseline (from a reference tip) over the entire run. This removes artifacts due to slight changes in buffer composition due to the increasing quantities of analyte as well as signal drift which is more common for non-streptavidin tips. Where multiple reference tips are used, they are averaged before subtraction. The second reference in double subtraction is the reference well. This involves performing a blank kinetic cycle using plain buffer during the experiment. To perform this subtraction, each kinetic cycle is first separated into their own trace. Each cycle begins with a baseline step, followed by an association step, and concluded with a dissociation step. These cycles are already primary reference subtracted with the reference tip, and now the blank cycle from each tip is subtracted from each individual kinetic cycle from the same tip. This helps correct for small variations in tip capacity and ligand loading density. Following double referencing, the cycles are then aligned to 0 by subtracting a constant value equal to the average of the last 5 seconds of the baseline step. As a final step, some software like the Octet Data Analysis HT perform a final data smoothing step, Savitzky-Golay filtering in this case. This is typically unnecessary but can help when signal-to-noise is low.

#### Models

It cannot be overstated that only 1:1 fitting should be relied upon for reliable kinetic parameters for BLI and SPR. Non-quantitative analysis of more complex binding is perfectly acceptable, but as we describe later, for *K*_D_ determination the more complex models are much more likely wrong than correct. The standard 1:1 fitting using the Langmuir-Hill equation ^43,44^, and is a well-established method that provides consistent results. This model was originally designed for modeling the adsorption of ideal gasses to solid surfaces, but it has found widespread use for biochemical kinetics. This involves fitting the reference-subtracted data to the integrated rate equations for the association (**Equation 5**) and dissociation phases (**Equation 6**). Where R_1_ is the signal observed during association at time t_1_ for analyte concentration [A], and R_2_ is the signal observed during dissociation at time t_2_.

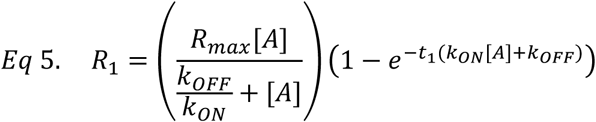

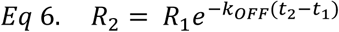

These equations can be used directly for fitting reference-subtracted data from a kinetics experiment, and they rely on several simplifying assumptions: 1) Ligands and analytes are homogeneous. 2) Each ligand binds a single analyte at an identical binding site with an identical binding energy. 3) Dynamic reversible equilibrium is established within the time-frame of the experiment. 4) No interaction between ligands or analytes should alter the binding at any other site. Clearly, these assumptions can never be strictly met, much like the assumptions required to use pseudo-first order kinetics in the first place, but for 1:1 binding without cooperativity the violations are not severe enough to warrant the use of a more complex model.

Issues with more complex models arise due to their increased degrees of freedom allow for improved fitting in many cases where the underlying biology does not support their use. For instance, a 2:1 heterogeneous ligand model and a 1:2 bivalent analyte model will often yield similar results regardless of the nature of the binding interaction. Worse than this, even when there is good justification to use a specific model, the limitations of current generation technology (both software and instrumentation) do not allow for accurate quantification. For example, we tested our nanobody saRBD-1 alongside bivalent Fc-saRBD-1, and then measured their *K*_D_s using both the standard 1:1, and the 1:2 bivalent analyte model. The results of these fittings are summarized in the table below (**Table 2**), but an important note is that this 1:2 model proports to give the *K*_D_ of the monovalent interaction by giving the *k*_ON_^1^ and *k*_OFF_^1^ equal to the true monovalent interaction while the *k*_ON_ and *k*_OFF_ give the amount of extra binding due to avidity. For both proteins, the fit improved, as shown by the Χ^#1^ and R^#1^ both dropping, but the *K*_D_ of the bivalent construct was not accurately recovered by using the 1:2 model. The accuracy of this method would be expected to increase with a lower affinity interaction, but the point remains that one cannot simply set up an experiment relying on this method to provide an accurate measurement; particularly when it is relatively easy to swap the bivalent molecule for the ligand to make an effective 1:1 interaction or digest the antibody to make a F(ab). Several kits (Fisher cat no. 44685) and well-established protocols^45^ exist for generating F(ab) molecules from traditional antibodies.

**Table 2:** Comparison of the 1:1 and 1:2 binding models using analytes of known stoichiometry.

| Protein | Model | $K_D$ | $k_{ON}^1$ | $k_{OFF}^1$ | $k_{ON}^2$ | $k_{OFF}^2$ | $X^2$ | $R^2$ |
| --- | --- | --- | --- | --- | --- | --- | --- | --- |
| saRBD-1 | 1:1 | 471pM | $3.36 \times 10^5$ | $1.58 \times 10^{-4}$ | N/A | N/A | 6.5378 | 0.9902 |
| Fc-saRBD-1 | 1:1 | <1pM | $1.57 \times 10^5$ | $<1 \times 10^{-7}$ | N/A | N/A | 53.5032 | 0.9899 |
| saRBD-1 | 1:2 | 893pM | $1.92 \times 10^5$ | $1.71 \times 10^{-4}$ | $1.31 \times 10^2$ | $2.32 \times 10^{-2}$ | 0.8553 | 0.9936 |
| Fc-saRBD-1 | 1:2 | <1pM | $5.00 \times 10^4$ | $<1 \times 10^{-7}$ | $2.98 \times 10^0$ | $1.15 \times 10^{-2}$ | 3.7041 | 0.9965 |

Using this data as an example again, the mass transport and heterogenous ligand binding models both also provide improved fits with reduced Χ^#1^ and R^#1^ values, showing that improved fits are generally to be expected any time a more complex model is used, even if inappropriately. This is not to say that it is impossible to use more complex models with BLI data, but one would need to know the stoichiometry prior to designing the experiment, carefully construct the experiment, and validate the findings using other approaches. A far simpler approach is to simply enforce 1:1 binding through experiment design.

For bivalent analytes, we already discussed either flipping the ligand/analyte, such that the monovalent molecule is now the analyte, or producing a monovalent version of the analyte either by digesting or inactivating one binding site. A second approach is to decrease the loading density until ligands are spaced too far apart for a single antibody to bridge the gap, but it can be difficult to tell when this has been achieved.

The mass transport model is designed to address the problem of analyte depletion in the local area of the sensor surface, but this is mainly seen in SPR due to laminar flow in the microfluidics whereas BLI uses turbulent flow to replace the surface layer much more quickly. Regardless, eliminating mass transport by decreasing loading density or increasing shake speed is preferable to using this analytical model. SPR studies and guides generally advocate for low density loading whenever possible as it reduces the likelihood of mass-transport limitation, and it reduces steric hinderance between analytes. For BLI, mass-transport limitation is unlikely due to the turbulent flow within the wells, but steric hinderance between analytes is a legitimate concern which should be considered.

The heterogenous ligand model assumes a mixture of two different binding sites in arbitrary proportions on the biosensor surface. This situation is theoretically possible, but unlikely. The more likely situation is that low quality ligand will contain a mixture functional, non-functional, and partially-functional binding sites which may lead to confusing results. This is best remedied by ensuring the quality of protein stocks prior to starting BLI.

In all cases, 1:1 binding is best suited for kinetics measurement, and lack of good fit should be an indication that more complex binding may be occurring. It is not good practice to observe which model fits best and assume that this describes the underlying biology. Instead, after seeing poor fitting in a 1:1 model, one should use alternative methods to determine which, if any, alternative models are appropriate, or if the protein quality and experimental parameters simply need to be optimized.

### Adaptation for non-VHH samples

This guide focuses on kinetic quantification and competition of VHHs, but the methods can be applied with few modifications to any number of other proteins. The primary caveat being that BLI is best for 1:1 interactions. Other immunoglobulin fragments like F(ab)s and scFvs can be tested in exactly the same way, while multivalent fragments like F(ab’)2 or intact immunoglobulins require modified protocols or digestion into monovalent fragments. For other non-immunoglobulins the stoichiometry may be less well-defined, and while BLI can still be a useful tool complex stoichiometry should be validated by other means (such as MALS) before claiming a *K*_D_. Advice on other individual proteins is highly case dependent, making it difficult to give broadly applicable guidance beyond the general good practice recommendations offered in this guide such as double referencing and trying both ligand/analyte orientations.

### Limitations

BLI is suitable for kinetic measurements of biomolecule pairs. This is possible as long as at least one of the components can be bound to a biosensor without damaging its activity. When possible, the larger component should be kept free in solution and the smaller component should be attached to the biosensor. We have experimentally verified good signal from proteins down to 11 kDa ^9^; we have anecdotally seen evidence of acceptable signal from 900 da molecules; and there is evidence from patents going down to 500 da ^46^. It is our belief that small molecules require exceptionally tight binding to display acceptable signal-to-noise in BLI. Competition assays can extend this lower limit further. On the upper end, BLI is capable of measuring whole bacteria as an analyte ^47^, and whole mammalian cells coated on tips ^48^, though signal in these assays does not yet outperform that of more conventional cellular assays.

Another limitation of BLI is that it cannot distinguish non-specific binding from specific binding, so this must be controlled experimentally. It also cannot measure the exact size of a complex, or precisely determine the residues involved in an interaction. However, residue-level information can be obtained through mutagenesis studies. While BLI is certainly capable of qualitatively measuring complex interactions, only 1:1 interactions should be used for kinetics calculations to determine *K*_D_, *k*_ON_, and *k*_OFF_. Currently, total experiment time is also a significant limitation of BLI due to evaporation issues, though this is primarily an engineering/protein stability challenge that will likely be solved if demand exists for 6+ hour experiments.

### Alternative approaches

Other methods to measure binding are many, though few are capable of providing the same depth of information as BLI. There are currently few BLI instrument manufacturers, with the Octet platform (currently owned by Sartorius) being the most popular. Gator Bio also produces a similar instrument. However, as discussed previously, SPR is nearly identical in terms of its data produced as well as its downstream processing. There are more instrument manufacturers of SPR systems including the popular Biacore platform (currently owned by Cytiva), and instruments produced by Nicoya (OpenSPR and Alto), Biosensing instrument (BI-4500), Bruker (Sierra platform). There are also hybrid systems such as the Fox Biosystems which use SPR measurement with a BLI-style fiberoptic sensor. In theory, data from any one of these instruments should be functionally equivalent, barring any platform-specific idiosyncrasies.

Many other methods of obtaining kinetics information, however most are much less user friendly. If labeling of both components is acceptable then Förster resonance energy transfer (FRET) is a good option ^49^, if only one component can be labeled then fluorescence anisotropy/polarization (FA/FP) is another possibility ^50^, and if labeling is not tolerated at all then classic radioligand studies or nuclear magnetic resonance (NMR) may be the only options. If one is looking for faster time scales than BLI can measure, stopped-flow experiments are a good choice that can be combined with any number of different detection methods including FRET, circular dichroism (CD), or simple fluorimetry ^51^. For epitope information one can do hydrogen-deutrerium exchange (HDX) ^10^, cryo-EM ^52^, or crystallography ^53^, though these can be difficult for non-experts. Additionally, computational modeling and simulation are increasingly used for predicting and analyzing these interactions.

For very precise, single-molecule work one might also try mechanical biosensors such as those used in atomic force microscopy (AFM), though these also require significant equipment and expertise ^54^. If equilibrium binding information is all that is needed, then other techniques such as isothermal calorimetry (ITC) ^55^, electrophoretic mobility shift assay (EMSA) ^56,57^, pull downs, or analytical ultracentrifugation are possible, but likely not easier than BLI. If a pseudo-*K*_D_ is acceptable then microscale thermophoresis (MST) ^58^ or thermal shift assay (TSA) ^59^ may possibly be faster depending on workflow, but *K*_D_ is often temperature dependent. One of the most recently developed technologies for measuring protein-protein interactions is mass photometry, which is a label free technique for measuring shifts in mass in the range of 20 kDa to 5 MDa ^60,61^. It is based on measuring the interference between the light scattered by the protein/protein-complex and the light reflected by the measurement surface. The signal measured is called the mass photometry contrast (or interferometric contrast) and is directly correlated with molecular mass. This technique is very powerful for determining equilibrium binding constants, but kinetic measurements of association and dissociation are more complicated than BLI. Lastly, EC_50_ values can be useful for early screening projects or when precise binding measurements are unimportant. Because of the broad definition of EC_50_, these can take the form of enzyme-linked immunosorbent assays (ELISA), functional enzyme assays, functional cellular assays, or any other test that can produce a dose-response curve. With the myriad of options available, it is worth strongly considering the upfront investment and ongoing cost associated with each in terms of money, time, and effort for the quality and applicability of data obtained.

## Materials

### Reagents

- Purified SARS-COV-2 RBD (BEI, Cat no. NR-52306)
- Purified Human Ace2 (Synbio, Cat no. 10108-H08B)
- Recombinant nanobody saRBD-1 (Fikadu Tafesse Cat# TafesseVHH_001, https://scicrunch.org/resolver/RRID:AB_3170195)
- Anti-SARS-CoV-2 Spike antibody B38 (InvivoGen Cat# cov2rbdc2-mab10, https://scicrunch.org/resolver/RRID:AB_3134136)
- Imdevimab (anti-SARS-CoV-2 Spike antibody) (InvivoGen Cat# srbdc4-mab10, https://scicrunch.org/resolver/RRID:AB_3134137)
- Bamlanivimab (anti-SARS-CoV-2 Spike antibody) (InvivoGen Cat# srbdc5-mab10, https://scicrunch.org/resolver/RRID:AB_3134138)
- MilliQ water
- HEPES (Fisher Scientific, Cat no. BP310-100)
- NaCl (Fisher Scientific, Cat no. S640-500)
- EDTA (Fisher Scientific, Cat no. S311-500)
- Tween-20 (Fisher Scientific, Cat no. AAJ20605AP)
- Bovine serum albumin (Fisher Scientific, Cat no. BP1600-100)
- Glycine (Fisher Scientific, Cat no. BP381-500)

### Equipment

- Octet RH16 (Sartorius, OCTET-RH16)

- Any Octet system will work with this protocol including previous ForteBio and Pall models.
- Streptavidin (SA) biosensor tips (Sartorius, Cat no. 18-5019)

- Tip chemistry is ligand dependent, see the section on Biosensors
- 384 well tilted bottom plates (Sartorius, Cat no. 18-5080)

- (Alternative) Greiner Bio-One black flat-bottom polypropylene

▪ 384 well (Greiner, Cat no. 781209)
▪ 96 well (Greiner, Cat no. 655209)

Caution – If using alternative plates, well volumes and biosensor height offsets must be adjusted. See note in procedure step 12.

- 1.5 mL Microcentrifuge tubes (Greiner, Cat no. 616281)
- Vortex mixer (Fisher Scientific, 14-955-151)
- Mini centrifuge (Corning, 6770)
- Micropipettes (0.1–2.5, 2–20, 20–200 and 100–1,000 µL; Eppendorf, Cat. nos. 3123000012, 3123000098, 3123000055 and 3123000063)
- Micropipette refill tips (10, 200 and 1,000 µL; STARLAB, Cat. nos. S1111-3700, S1111-0706-C and S1111-6701-C)

### Software

- Octet Data Acquisition (Sartorius)
- Octet Data Analysis HT (Sartorius)
- Example method files (included in Supplemental information)

### Reagent Setup

#### Biotinylated ligand protein stock

Ligand protein should be biotinylated for use with streptavidin biosensors. A labeling efficiency of 1-2 biotins/protein is ideal. Box 1 shows a protocol for biotinylation of a protein of interest. The protein should be between 0.5 - 10 mg/mL in sample buffer and should be thawed on the same day as the experiment.

#### Analyte protein stock

Analyte protein should not contain biotin, to avoid interaction with the biosensor surface. The protein stock should be between 0.5 – 10 mg/mL in sample buffer and should be thawed on the same day as the experiment.

Critical – The quality and purity of the analyte protein is essential to obtaining good results. Also, accurate information about molecular weight and concentration are key as they are directly input into the analysis equations.

#### Sample buffer

Starting with MilliQ water, add HEPES to 10mM and adjust to pH to 7.5. Add NaCl to 150mM and EDTA to 3mM. Store at room temperature and use within one month.

Optionally, use commercial Octet Sample Diluent (Sartorius, Cat no. 18-1104). Alternative sample buffers can be used. See section on the Blocking step under Experimental design.

#### Running buffer

Starting with sample buffer, add 0.005% (v/v) Tween-20 and 0.1% (w/v) BSA. Store at room temperature and use within one week.

Optionally, use commercial Octet 10x kinetics buffer (Sartorius, Cat no. 18-1105).

Alternative running buffers can be used. See section on the Blocking step under Experimental design.

#### Regeneration buffer

Starting with MilliQ water, add Glycine to 10mM and adjust to pH 1.7. Store at room temperature and use within 6 months.

Alternative regeneration buffers can be used. See section on the Regeneration step under Experimental design.

### Procedure 1: Basic kinetics

Timing 3 hours

Critical – This protocol assumes access to a set up an properly maintained Octet system with paired computer set up with Octet Data Acquisition software.

#### Stage I: Experiment design

1) Open the Octet Data Acquisition software

a) The Octet machine will take ∼1 min to start up
2) In the experiment wizard, select basic kinetics or load a previous method file (see example method file in the Supplemental Information).

[See Troubleshooting]

3) Enter the plate design in tab 1 (**Figure 4)**.

a) Plates 1 and 2 can both be used if more samples are needed.
b) Proper Analyte concentration is critical at this step because the value entered at this step will be carried forward into the data analysis and cannot be easily changed later.

**Figure 4:**
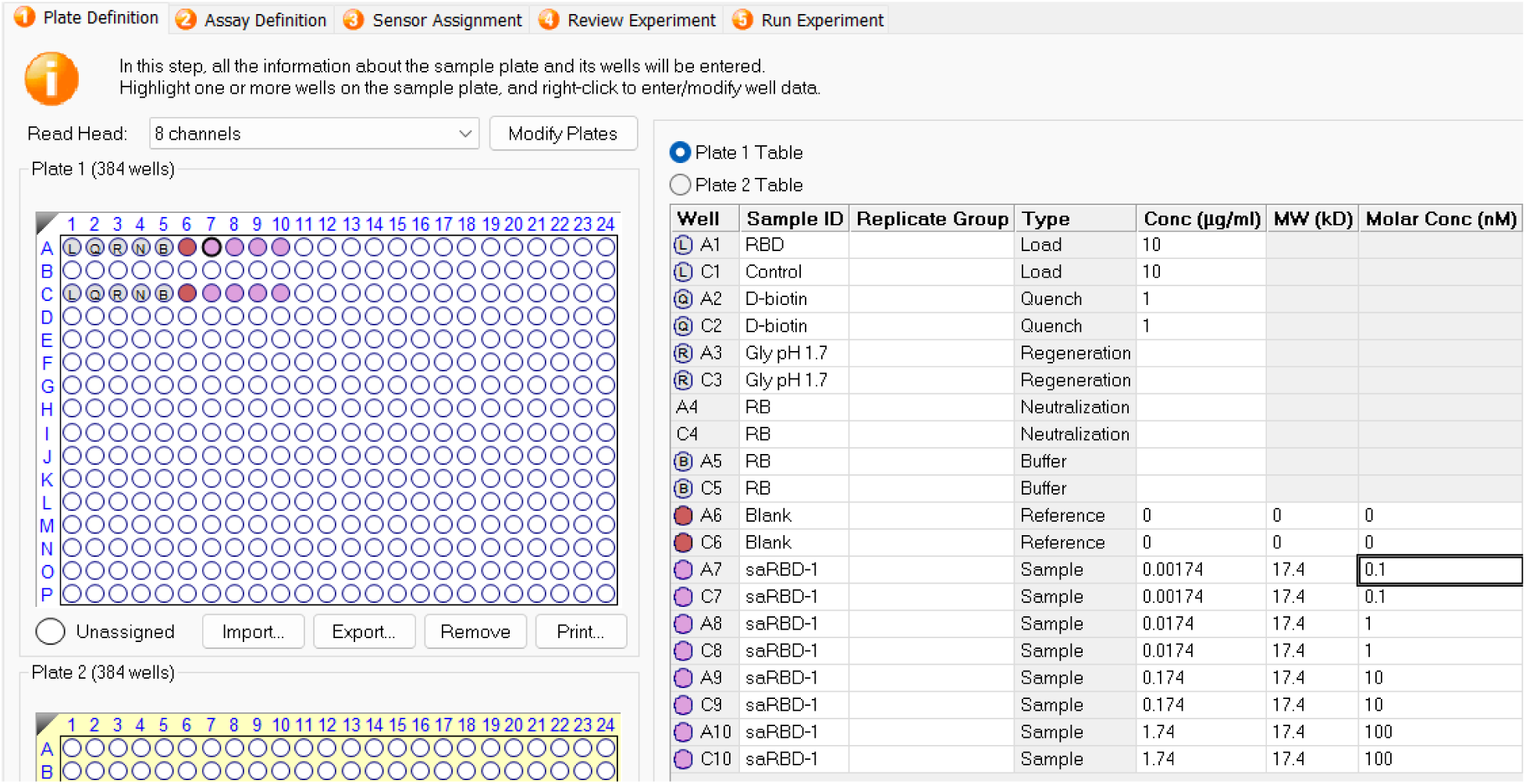
Plate layout tab with example data.

Critical step – Samples intended for the same tip must be in the same row, or one below. For example, tip 1 can enter wells A1-A24 and B1-B24 while tip 2 can only enter wells C1-C24 and D1-D24.

Critical step – Some step types are not optional as they can affect the interpretation of the data by the analysis software. The step types are defined as follows:

- Sample – Analyte sample, these wells are for association steps only

- Data may not be recognized properly by analysis software if this step is not used, or used excessively for other purposes.
- Reference – Plain buffer for background subtraction, this can be assigned later during analysis
- Control – Known samples for confirming proper assay behavior, not used for analysis
- Buffer – Similar to reference, plain buffer for any purpose
- Activation – only used for chemically activated tips like amine-reactive
- Quench – generally to quench chemical reactions, also useful for biotin blocking streptavidin
- Loading – ligand to load on the biosensors
- Wash – similar to buffer
- Regeneration – specifically for regeneration steps, must contain regeneration buffer
- Neutralization – Must be present if a regeneration step is used, generally contains RB

4) Enter the step definitions in tab 2.

a) Sample step definitions (**Figure 5**).
b) Sample regeneration parameters in (**Figure 6**).
c) To add a step, click “Add” under “Step Data List”, see **(Figure 7)**.

i) In this example, a step type called “baseline” which lasts 30 seconds is being added
ii) There is no need to adjust rpm
iii) Regeneration steps cannot be modified here, first add to the main list, then select “regeneration params” and change regeneration cycle times and counts in the new window.
d) To start a new assay:

i) Select a step type – it will be highlighted (**Figure 5)**.
ii) Click on the well that step will take place in (must be rows A or B) – all 8 (or 16) selected wells will be highlighted (**Figure 5**).
iii) Click “New Assay”
e) To add a step to the assay:

i) Select a step type
ii) Double click the well that step should dip into
iii) For Regeneration steps, click edit step to confirm correct assignment of neutralization wells

**Figure 5:**
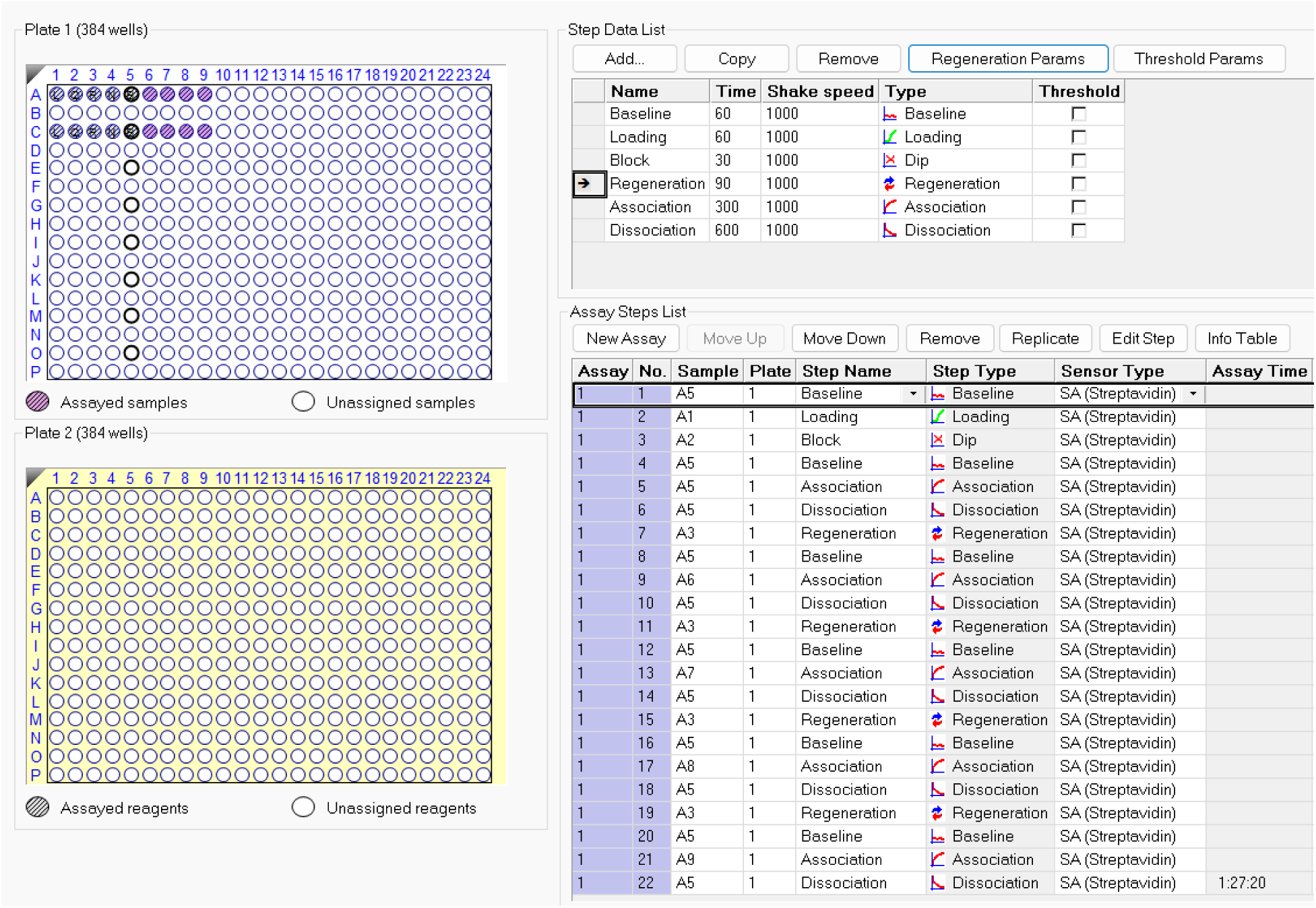
Step definition tab with example data.

**Figure 6:**
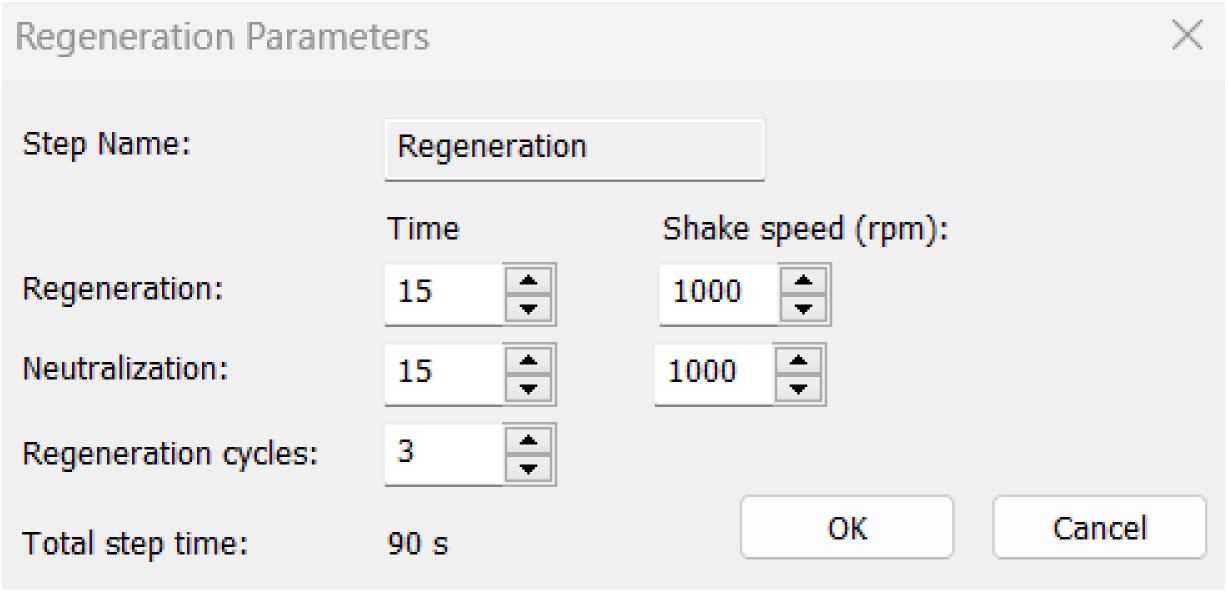
Regeneration parameters with example values.

**Figure 7:**
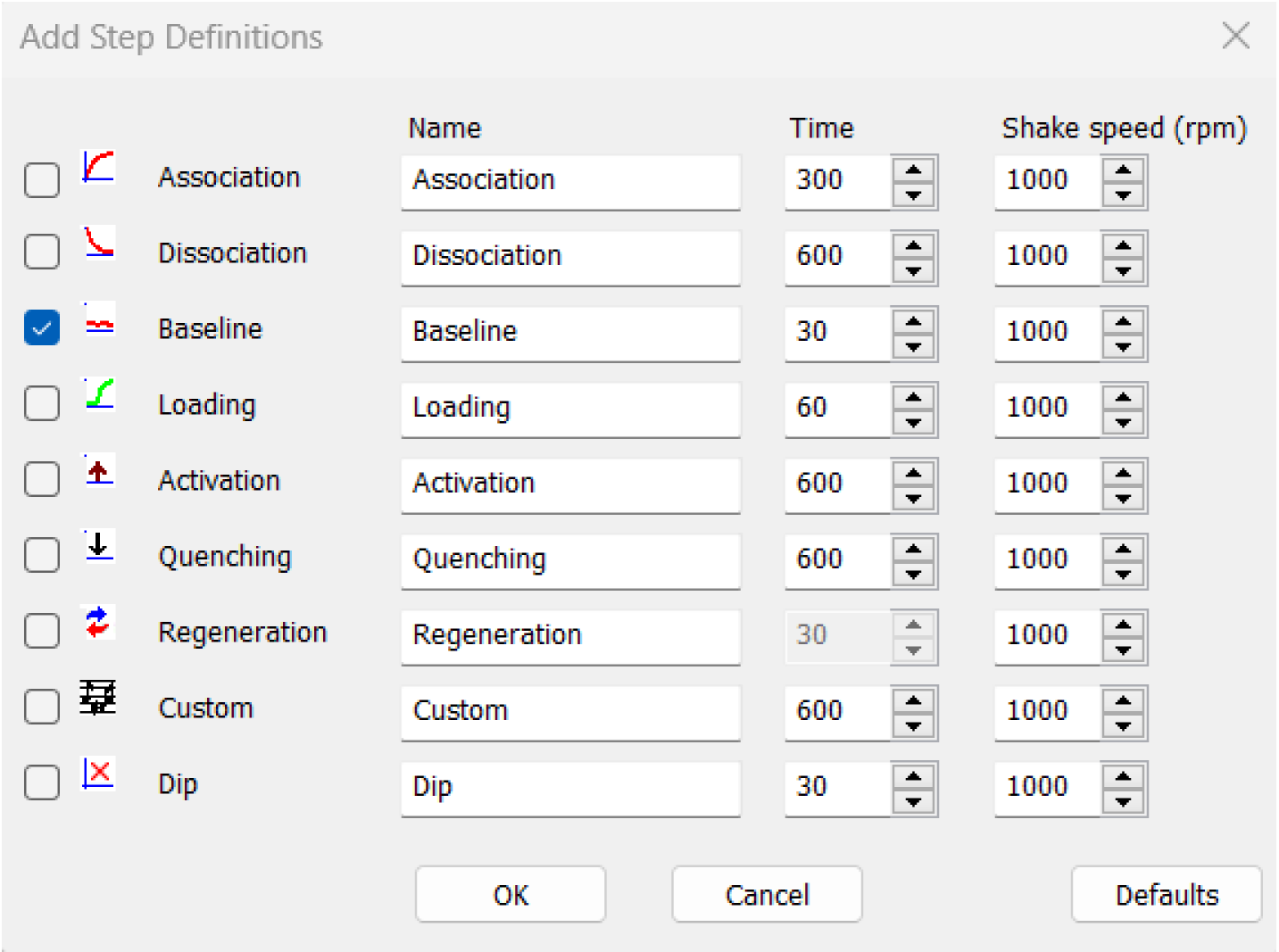
Add step definitions menu. The times listed are a good starting point, but should be adjusted to meet the needs of each individual experiment (see **Tables 3 and 4**).

**Table 3:** Equilibrium concentrations.

| 300 second association | | $k_{OFF}$ | | | |
| --- | --- | --- | --- | --- | --- |
| | | $10^{-2} s^{-1}$ | $10^{-3} s^{-1}$ | $10^{-4} s^{-1}$ | $10^{-5} s^{-1}$ |
| $k_{ON}$ | $10^4 M^{-1}s^{-1}$ | 500 nM | 1400 nM | 1500 nM | 1500 nM |
| | $10^5 M^{-1}s^{-1}$ | 50 nM | 140 nM | 150 nM | 150 nM |
| | $10^6 M^{-1}s^{-1}$ | 6 nM | 13 nM | 15 nM | 15 nM |

**Table 4:** Dissociation step times required to reach 5% signal reduction.

| $k_{OFF}$ | $10^{-3} s^{-1}$ | $5 \times 10^{-4} s^{-1}$ | $10^{-4} s^{-1}$ | $5 \times 10^{-5} s^{-1}$ | $10^{-5} s^{-1}$ |
| --- | --- | --- | --- | --- | --- |
| time to reach 5% dissociation | 50 s | 100 s | 500 s | 1000 s | 5000 s |

Critical step – The total experiment should not exceed 3 hours (2 preferred) to prevent issues with evaporation. Samples will concentrate over time as evaporation occurs and the fluid level in the well will eventually drop too low for the sensor to reach, also increasing protein concentration.

5) Sensor assignment is automatic and happens in tab 3 (**Figure 8)**.

a) Blue squares indicate these sensors will be used in the assay
b) The Octet will only pick up tips from the blue locations during the experiment

i) If no squares are blue, double check that the sensor type assigned in the assay matches the type assigned to the plate.
ii) Right click on a square to assign a different sensor type.
iii) Lot can be used to assign replicates during analysis.
6) Review the experiment in tab 4 by stepping through each step and ensure that the correct step type is occurring in the correct order and in the correct well.
7) Run the experiment (**Figure 9).**

a) Assign file names as appropriate
b) Delayed start is useful if the biosensors have not been pre-soaked prior to the start of the experiment, but is otherwise unnecessary
c) Ensure the temperature is at the expected 30°C unless otherwise specified
d) Sensor offset should generally be set to 4, except for tilted well plates which should be 3.

i) Adjusting this can change when evaporation becomes a critical concern, and how much reflected light from the well bottom re-enters the tip, adding noise.
e) Acquisition rate does not change the underlying measurement frequency, just the number of measurements that are averaged into each value.
8) Ensure that the sample plate(s) is prepared and placed in the proper location in the Octet, and that the biosensor soaking plate and rack are properly installed with the correct biosensors, prior to starting the experiment by clicking “Go”

**Figure 8:**
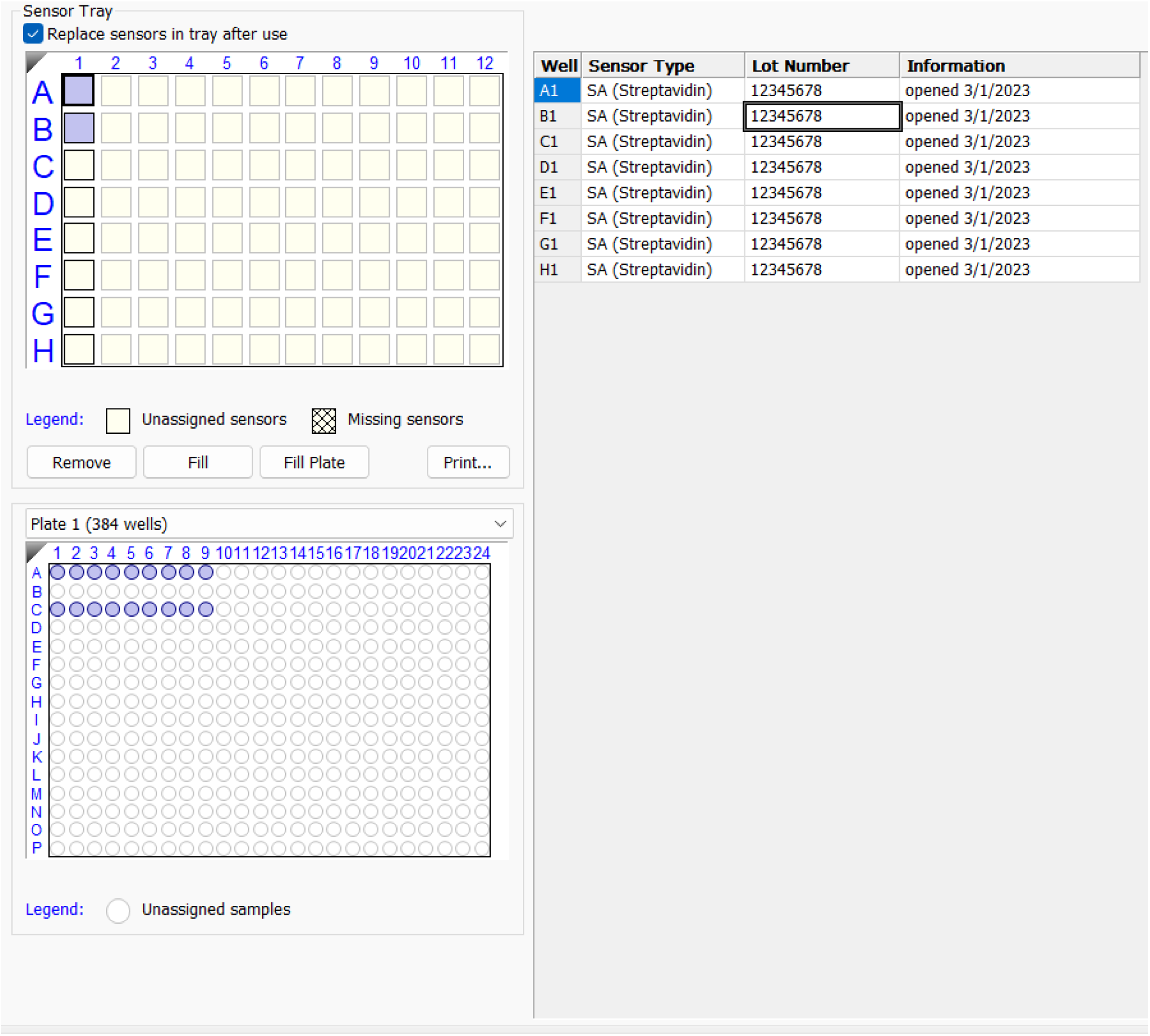
Sensor Selection.

**Figure 9:**
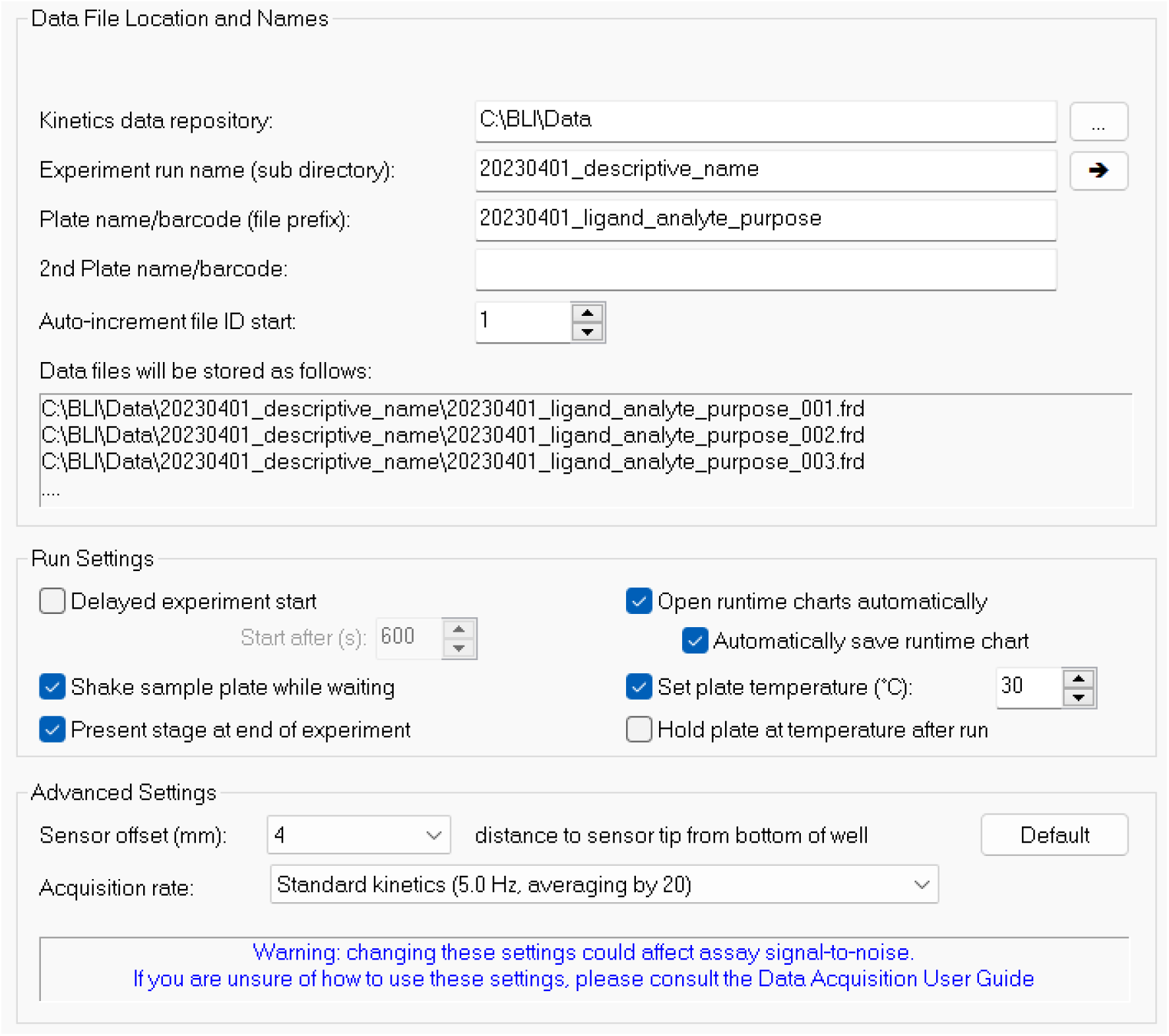
Pre-run settings.

Pause point – The experimental design can be created prior to the time of the experiment and saved as a (.fmf) method file. Open the method file with the octet software, and confirm design and run parameters before starting the run.

#### Stage II: Run the experiment

9) Prepare the following samples in 1.5mL microcentrifuge tubes:

a) >1mL MilliQ water
b) >1mL Running buffer (RB)
c) >1mL Regeneration buffer
d) 120µL Ligand, 10µg/mL in RB
e) 150uL Analyte, serially diluted in RB in the following concentrations:

i) 100 nM
ii) 10 nM
iii) 1 nM
vi) 0.1 nM
10) Vortex and spin samples down samples, then store at room temperature until use

a) Use within 12 hours or less depending on protein stability.
11) Prepare the biosensors

a) Load a 96 well plate into the biosensor slot in the Octet
b) Fill the wells with ∼150µL of MilliQ water
c) Place an empty biosensor tip rack over the soaking plate
d) Carefully place the biosensors into the rack so that they dip into the liquid below.

i) The tip of each biosensor much not touch any surface when removing from the packaging and placing in the Octet machine.
ii) Use a 10µL pipette (single- or multi-channel) to transfer tips
e) Soak for a minimum of 20 minutes prior to starting the experiment, use this time to prepare the sample plate and review the experiment design.

[See Troubleshooting]

12) Prepare sample plate according to the design from step 3.

a) Sample volumes differ between plates

i) Tilted well 384 – 45-60 µL
ii) Flat bottom 384 well – 80-120 µL
iii) Flat bottom 96 well – 150-200 µL
b) Adjust sensor offset accordingly in tab 5.
c) Place the plate in the correct location and orientation of the Octet.

i) Empty plates need not be placed in their slots
13) Review method and press “Go”
14) A new window will open with information about the run, and limited controls for modifying it (**Figure 10**).

a) The only controls after starting the run are:

i) Stop the run
ii) Extend the current step – requests a number of seconds to extend the step by.
iii) Go to next step – moves on to the next step immediately.
iv) These can only modify a step that is already in process, not future steps.
v) These are mostly useful for manually adjusting the loading times. [See Troubleshooting]
15) Let the run complete and collect the data from the created folder.
16) Remove samples and biosensors from the Octet and close the lid.

**Figure 10:**
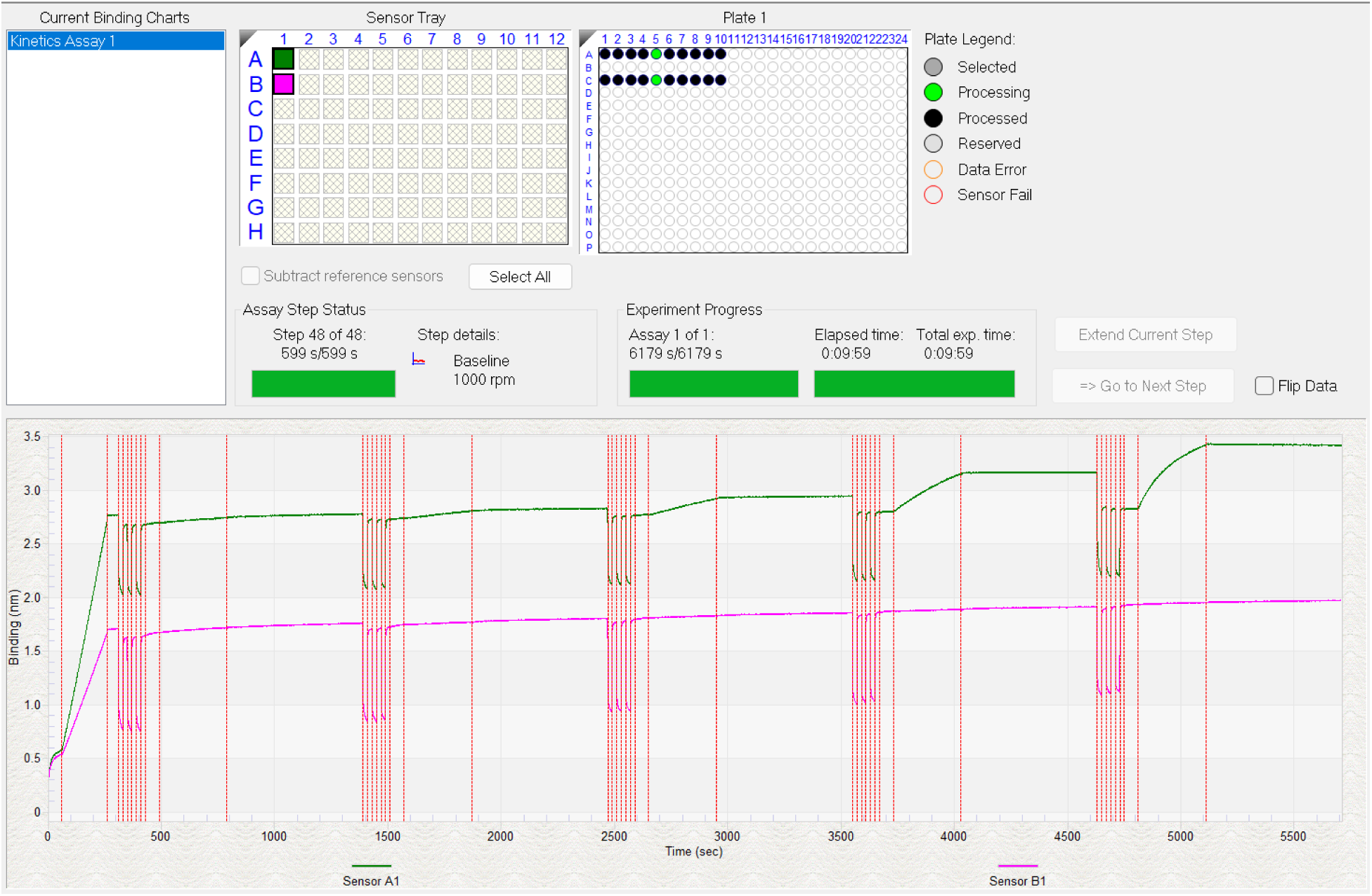
Example runtime display based on the above experiment design. Extend step and go to next step buttons are greyed out because the run is complete.

Pause point – Once the data is properly saved analysis can happen later on any computer that has the appropriate software.

#### Stage III: Data analysis

17) Open the Data Analysis HT software

a) The standard Data Analysis software has fewer features, so is not very useful.
18) Open the data file by navigating to it in the left hand box and double clicking on it.
19) Once open, click on pre-process operations in the top bar (**Figure 11)**
20) If not assigned during experiment setup, designate reference sensors.

a) Select all intended reference sensors with Control + left mouse button
b) Selected sensors will have a blue border
c) Right click and select reference sensor in the sensor type menu
21) Perform primary reference subtraction

a) Select all sensors with Control + left mouse button
b) Right click and select “subtract reference in selected well”
c) This will average all selected reference sensors and subtract from all standard sensors
d) If some sensors have specific reference sensors, select each group separately for step b
22)Enter the “Reference Sample” tab (**Figure 12)**
23) Select wells used for double referencing

a) Hover over wells to see their contents
b) Selecting a well will also automatically select the buffer well used for dissociation
c) Right click and set to reference sample type similar to step 20
24) Perform double reference subtraction

a) Select one reference well
b) Using control + left mouse button, select all sample wells associated with this reference
c) Right click and select “subtract reference in selected wells
d) Perform steps a-c for each reference
e) More than one reference can be used, and will be averaged

i) Sample references should generally be used internally within each biosensor
25) Confirm proper ref subtraction and double ref subtraction formulas in the sample list, and confirm simple exponential curve shapes with increasing curve steepness and R_max_.

[See Troubleshooting]

26) Enter the “Data Correction” tab
27) Align Y Axis

a) Set to “average of baseline step”
b) The default, 5 second window at the end of the baseline step, is usually good
28) Inter-step correction should not generally be used

a) This will partially cancel large buffer spikes, but it is always superior to use proper reference subtraction.
29) Leave the box for Savitzky-Golay Filtering checked

a) This is a smoothing function similar to a rolling average
b) No substantial effect on fitting is observed in most cases
30) Select the “Kinetic” button in the top right of the window (**Figure 13)**
31) Steps to analyze

a) Set to “Association and Dissociation”
32) Binding model

a) Set to “1:1”
b) The 1:1 model is the only model that can reliably determine binding kinetics parameters.

i) See section on binding models for further discussion of other models
33) Fitting

a) Set fitting to “Local”

i) Local fits are best for pilot experiments where many concentrations will not be within the ideal range.
b) Set Fit steps to “Full”
34) Window of interest should not generally be modified
35) Steady state analysis can only be used for molecules with fast kinetics that reach saturation at every concentration.
36) Use shift/control + left mouse button in the sample list on the lower right side, then select individual curves to examine fit lines
37) Fitting parameters (*K*_D_, *k*_ON_, *k*_OFF_, R_max_, *k*_obs_, errors, statistics, etc.) are on the right side of the sample list.
38) Find the concentration with the most even curvature (**Figure 14**)

a) this will likely also have the closest fit

**Figure 11:**
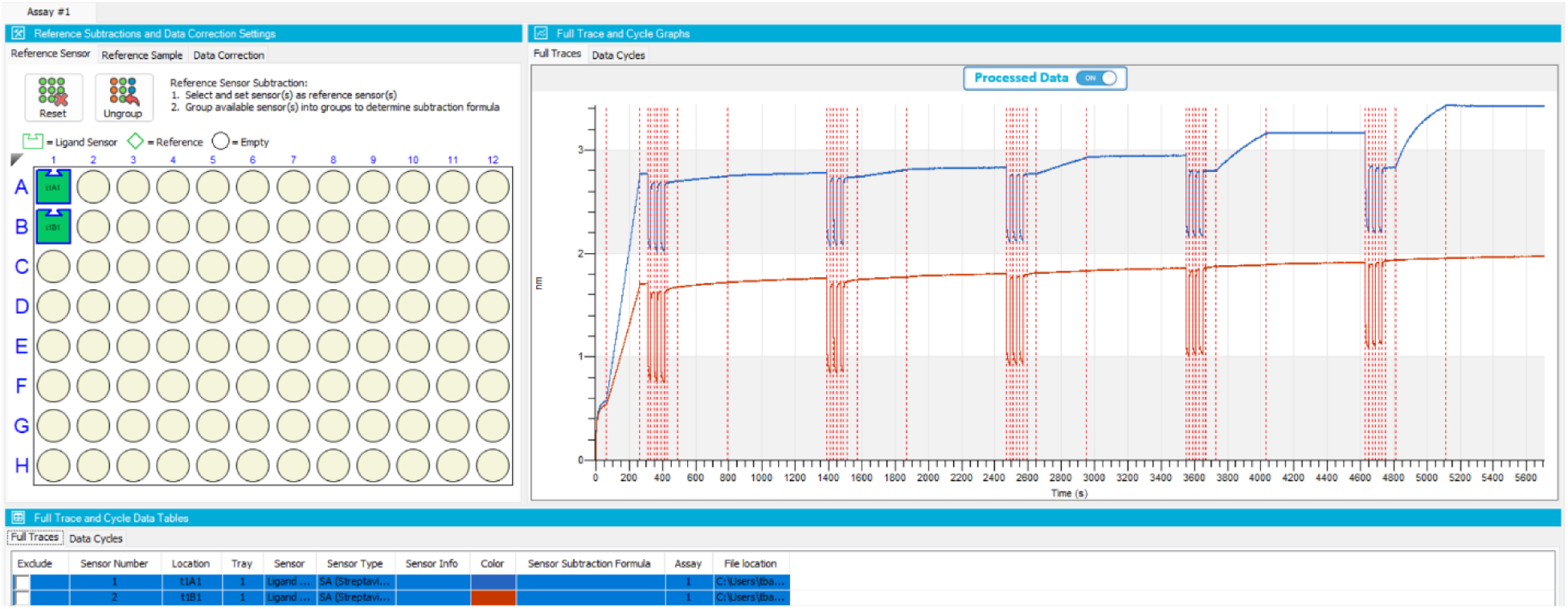
Reference sensor selection tab with example data.

**Figure 12:**
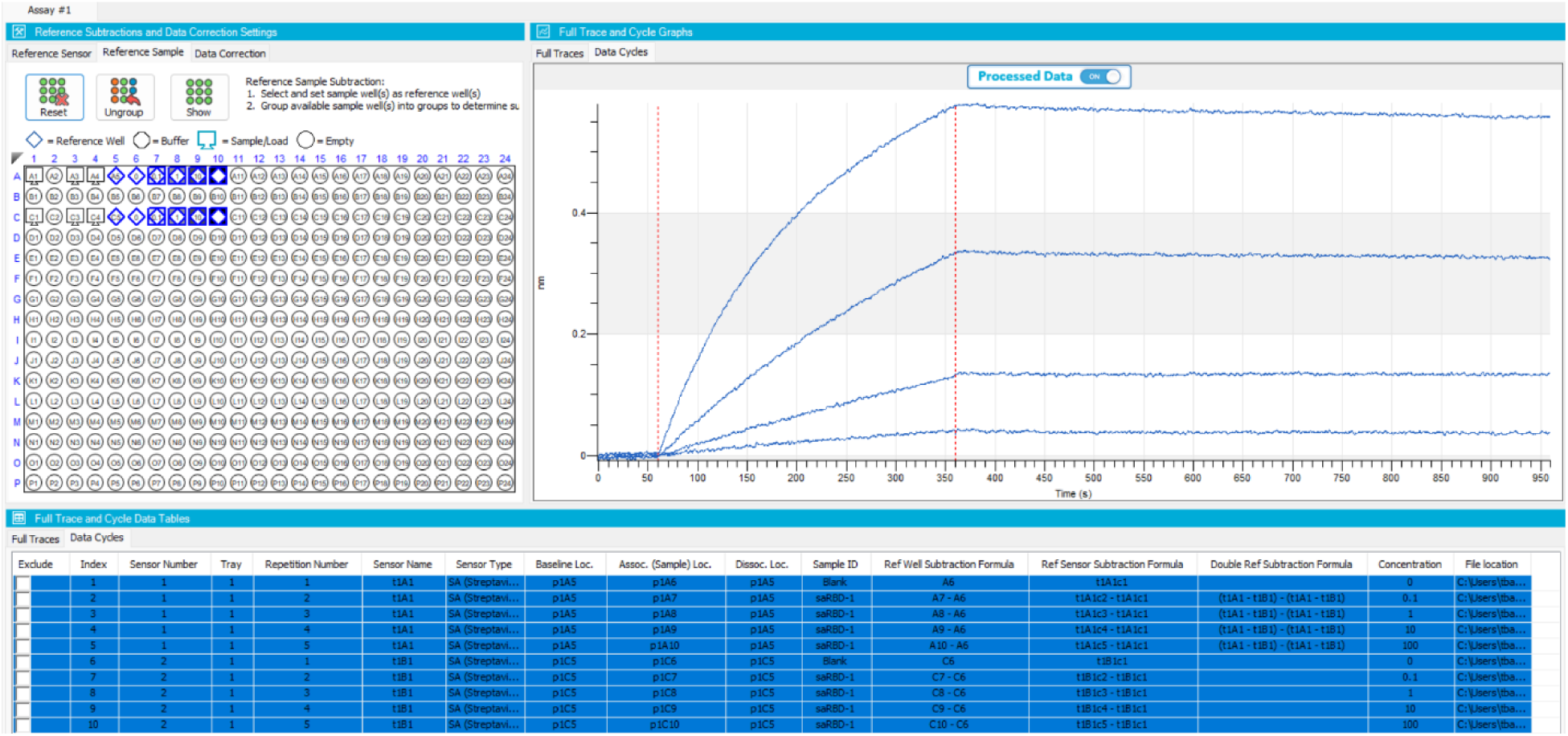
Reference Sample selection with example data.

**Figure 13:**
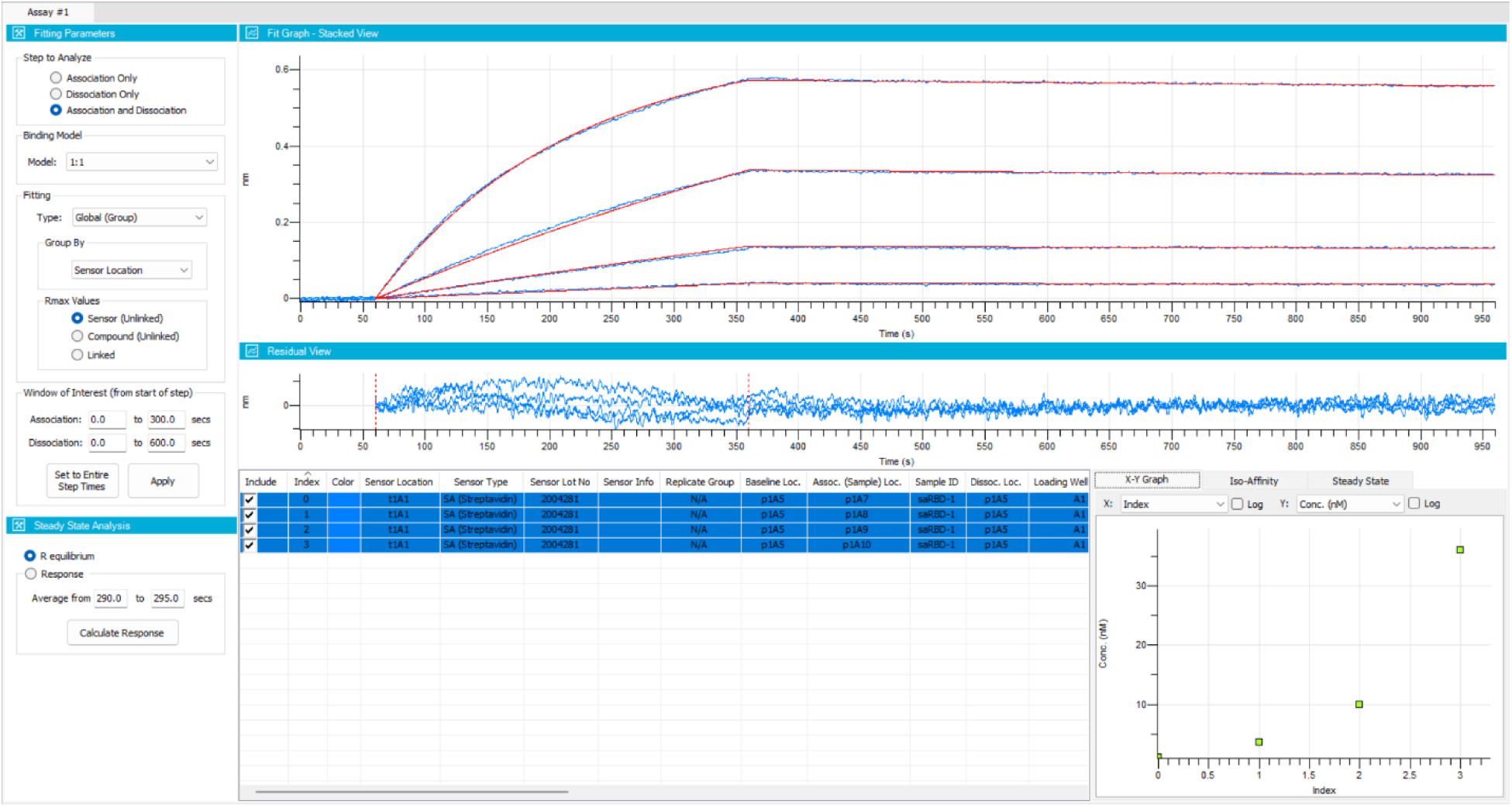
Kinetic fitting tab with example data.

**Figure 14:**
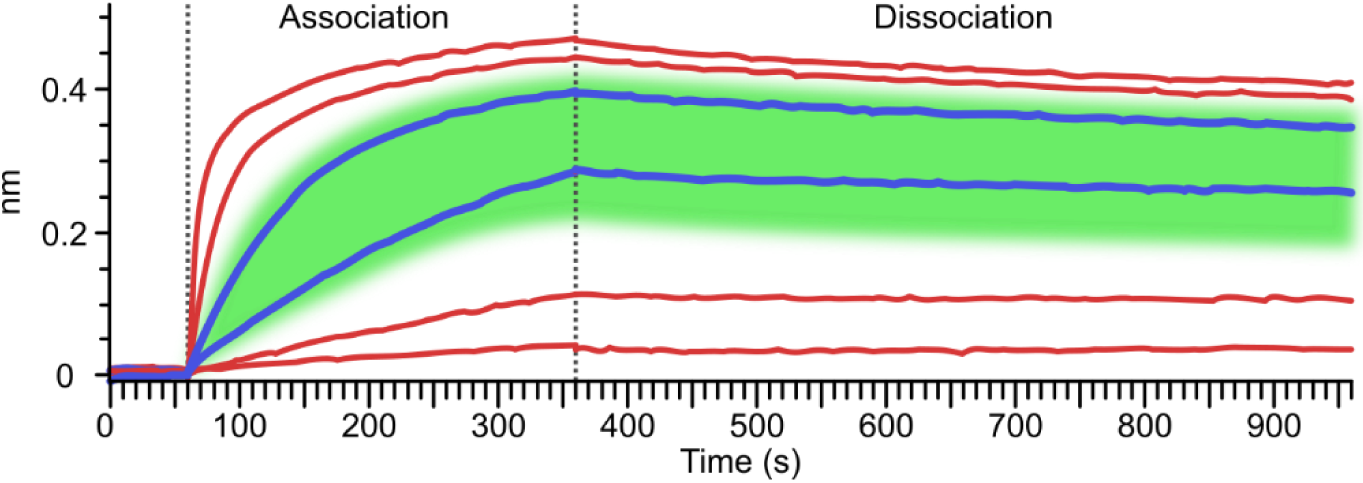
Selecting the ideal curve for *K*_D_ estimation from a pilot experiment. The green area shows the approximate ideal range for most accurate *K*_D_ estimation using local fitting. The blue curves should be selected in this case as they fall within this range. Lower concentrations do not have enough curvature, while higher concentrations are more likely to have biphasic curves due to non-specific binding. Note that concentrations here are half-log (1:3.16) dilutions, and most 1:10 dilution schemes will only have one curve in or near the ideal range.

[See Troubleshooting]

33) Observe the *K*_D_ calculated for the selected concentrations. This will be the estimated *K*_D_ that will be used to design the full kinetics experiment.

### Procedure 2: Full kinetics experiment

Timing 3 hours

1) Perform pilot experiments with analyte to estimate *K*_D_.
2) Start a new method using the basic kinetics template in the wizard or start with a previous kinetics assay method (see example method file in the Supplemental Information).
3) Design an experiment following the steps of Procedure 1 steps 1-8 with the following changes (see example plate setup in **Figure 15**).

a) Include 2-3 replicates of each ligand-analyte pair

i) Also include 2-3 replicates of control tips
b) Include 5-7 concentrations with 2-3 fold dilutions.

i) Initially, use the ideal concentration from the pilot experiment at the center of the dilution series.
Start with a similar step definition and order as the pilot experiment.
Adjust the loading sample concentration based on observed signal in the pilot experiment

a. Target 1 nm of loading binding for all load samples and controls (except empty)
b. For equal signal, calculate loading relative to load protein and analyte sizes

i) See section on loading for more details.
Adjust the association step time

a. An association of 300 seconds is typical.

i) For weak interactions, association time can be decreased.
ii) For strong interactions, concentration can be increased rather than time.
b. **Equation 3** can be used to calculate time to reach equilibrium, and (**Table 3**) shows the concentrations necessary to reach 99% equilibrium for various combinations of *k*_ON_/*k*_OFF_ values within 300 seconds.
c. Alternatively, use an SPR/BLI sensorgram simulator to estimate appropriate concentration ranges and step lengths (e.g. https://apps.cytivalifesciences.com/spr/)
Adjust the dissociation time so that at least 5% signal reduction will occur.

a. For a 5-concentration assay, 600 seconds is a good dissociation step time for tight binders (*k*_OFF_ ≈ 10^-4^ s^-1^).
b. For even tighter binders, shorten dissociation time to 60 seconds for all except the highest concentration, which should be increased to 3,000 seconds.

i) Double referencing will not work with this method, so use reference tips only.
ii) Alternatively, have separate reference steps for both dissociation step lengths, but this will make the experiment much longer.
c. This time can be calculated using **Equation 4**, and (**Table 4**) shows times needed to reach 5% dissociation for various *k_OFF_* rates.
Reassess the final experimental timing and add or remove concentrations in order to keep the total time around 2 hours.

a. A 6-concentration experiment (not including reference step) with 300 second association and 600 second dissociation steps will take around 7800 seconds, or just over 2 hours.
Run as normal, following Procedure 1 steps 9-16.
Analyze following Procedure 1 from step 17 up until step 33 (fitting).
Fitting

a. Set to “Global”
b. Group by “color” is generally useful.

i) Colors can be set automatically or manually at a later step.
c. Alternatively, group by a different parameter that adequately organizes the replicates.
R_max_ Values

a. Set to “Sensor (unlinked)”
b. Linking forces all samples to use the same R_max_ value which is related to sensor capacity

i) Sensor (unlinked) assumes the R_max_ will be the same only when on the same sensor
ii) This behaves the same as linked when all grouped samples are on the same sensor
Observe the final fit, does it meet these criteria?

a. There are at least 3 concentrations with strong curvature

i) Not a flat line or linear increase (low concentration)
ii) Does not reach saturation in under 5-10 seconds (high concentration)
b. The fit lines accurately match the data for all concentrations

i) Observe the residual trace for any large variations [See Troubleshooting]
Some samples may have excessively low or high binding or other artefacts. These samples can be excluded from the analysis by unchecking the “include” box

a. Often High concentration samples have non-standard binding due to concentration-dependent non-specific binding (**Figure 16**).
b. Often very low concentration samples will display less signal than expected.
c. Other possible artefacts are:

i) sudden jumps in signal mid-step due to biosensor damage
ii) Rapid and large spikes of 10-20 nm due to air bubbles or loss of biosensor contact with the liquid surface.
Confirm results by performing a second repeat assay on a different day, ideally with different batches of ligand and analyte.

**Figure 15:**
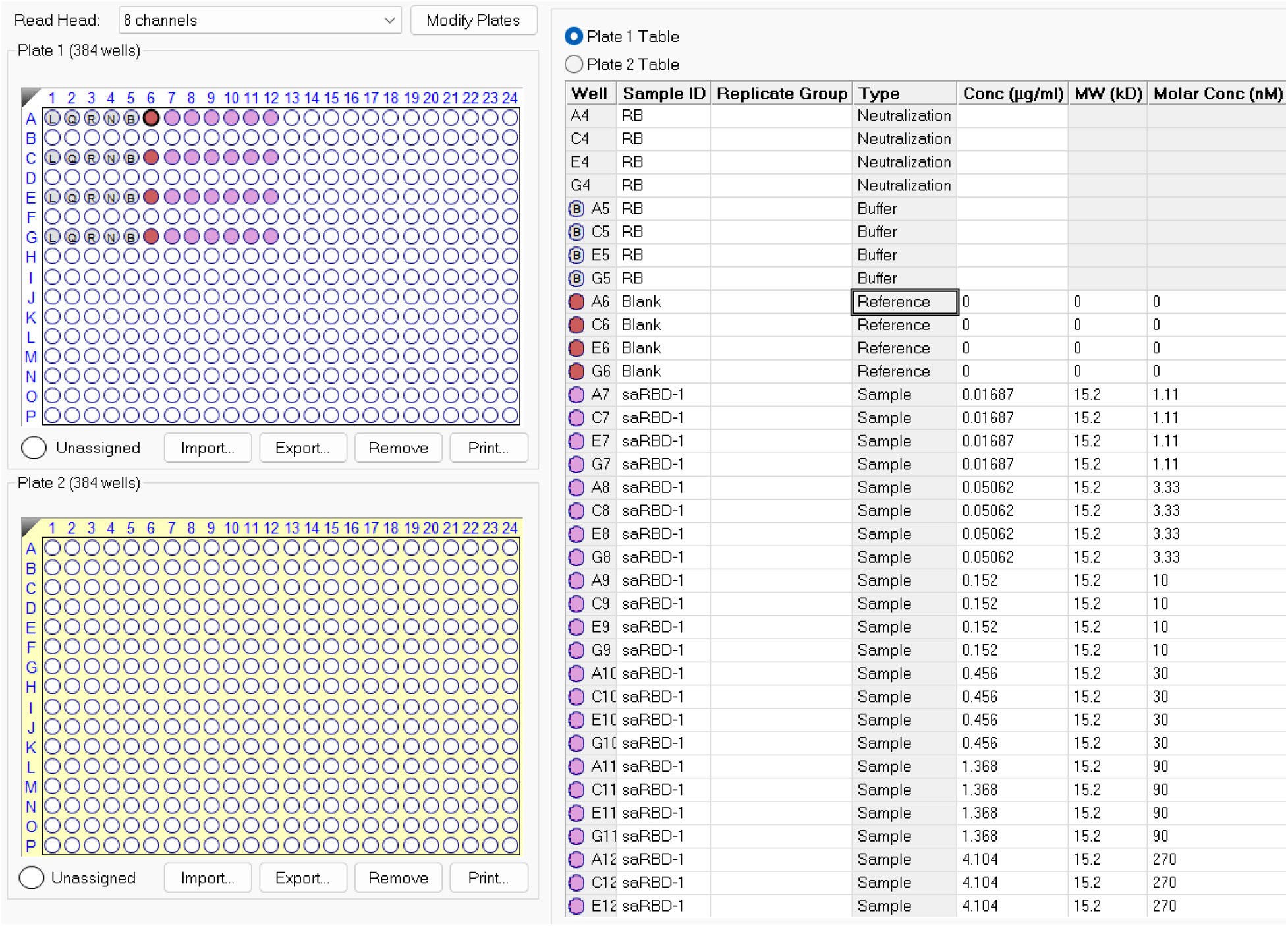
Example plate layout for a full kinetics experiment.

**Figure 16:**
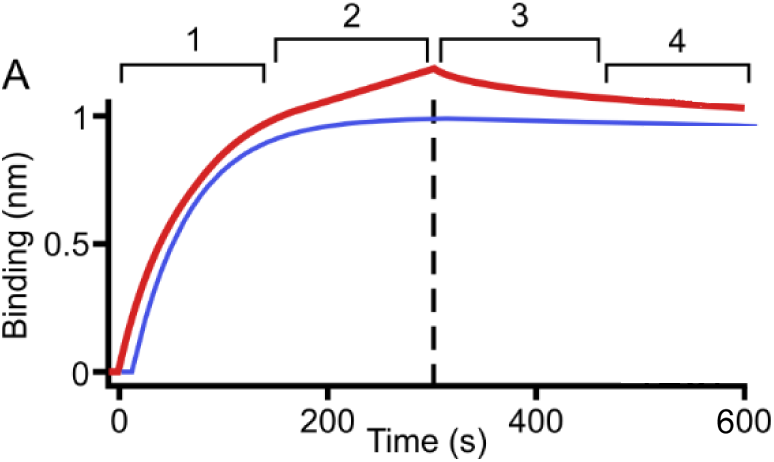
Non-standard curve shape due to concentration-dependent non-specific binding. In blue is a perfect exponential association and dissociation (albeit slow). In red is a sample with non-ideal binding at this concentration. Stages 1-4 show different aspects of this. (1) Apparently normal binding that follows an exponential. (2) A continued, slower, often linear increase due to non-specific binding. (3) An initial rapid but small drop at the beginning of dissociation. (4) return to expected dissociation curve. Samples exhibiting this behavior may or may not eventually return to 0 nm of signal.

### Procedure 3: Competition assay

Timing 3 hours

1) Perform pilot experiments with analyte and competitor to estimate *K*_D_ for each.
2) Start a new method using the basic kinetics template in the wizard or start with a previous competition assay method (see example method file in the Supplemental Information).
3) Design an experiment following the steps of Procedure 1 with the following changes (see example plate setup in **Figure 17**).

a)Include 2-3 replicates of each ligand-analyte pair

i) Also include 2-3 replicates of control tips
b) Include 5-7 concentrations with 2-3 fold dilutions.

i) Ensure that the highest analyte concentration is near saturating

**Figure 17:**
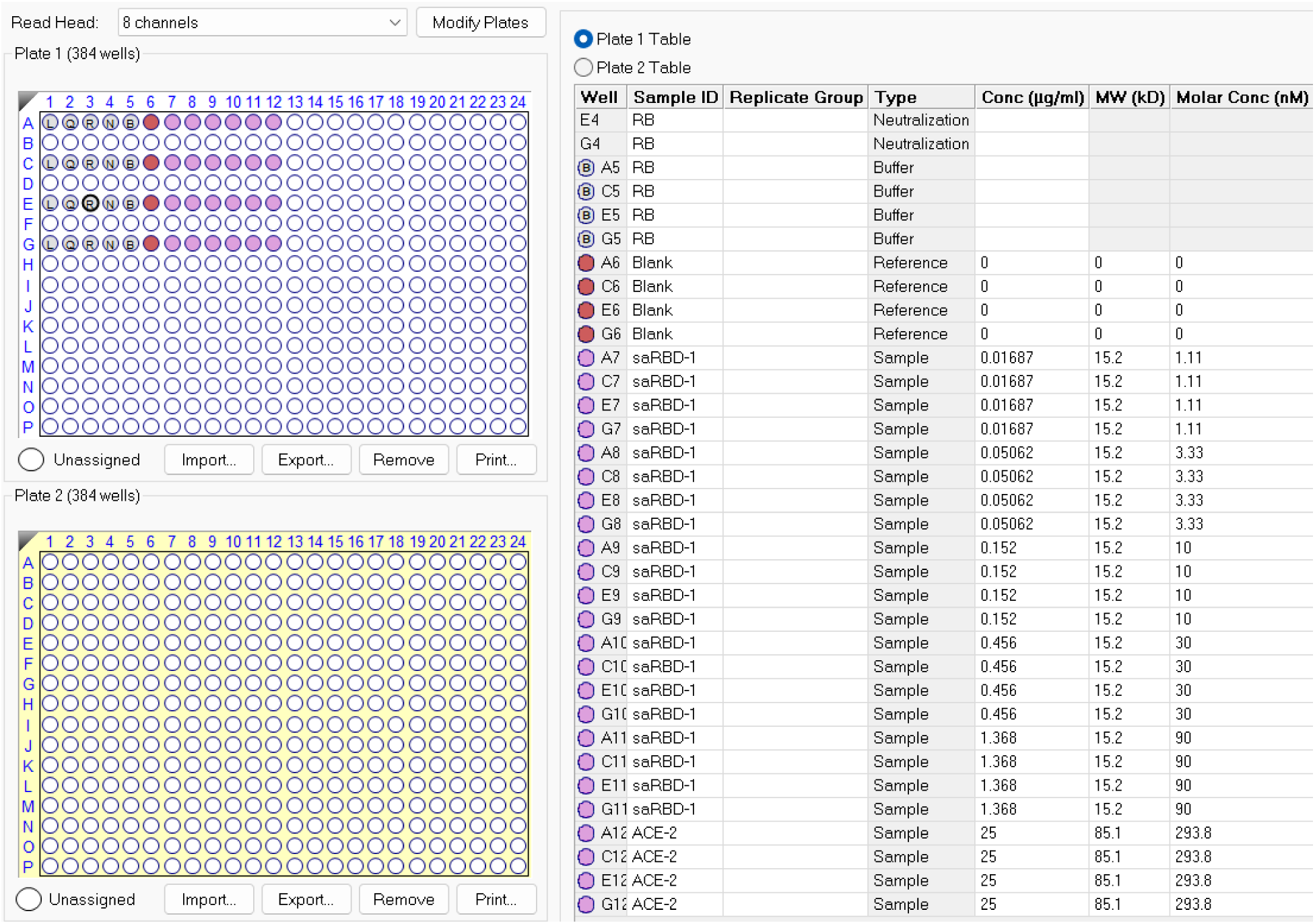
Example plate layout for a competition experiment.

Critical step – The only way to definitively tell if a concentration is near saturation is to plot multiple concentrations on a dose-response curve. Every concentration will reach steady-state equilibrium and level out, and higher concentrations will asymptotically approach saturation (**Figure 18**). A shortcut approach is to test escalating concentrations.

c) Include a competitor well with high enough concentration to get strong binding within a short time, without causing non-specific binding.

4) Start with a similar step definition and order as the pilot experiment.
5) Set up the association times to be the shortest times that allow for appropriate saturation based on pilot experiments.

a) Primary analyte and competitor analyte do not need to use the same timing.
6) No Dissociation times are needed for competition experiments.

a) Make note of the dissociation rate of the primary analyte.
b) This protocol works best when minimal (<5%) dissociation of the primary analyte occurs over the course of competitive analyte association steps.
c) It is possible but not trivial to correct for primary analyte dissociation.
7) The final experiment should follow a similar pattern to the example in (**Figure 19**).
8) Run as normal, following Procedure 1 steps 9-16.
9) Analyze following Procedure 1 steps 17 through 30 (kinetics tab)
10) Right click on the sample list area and click “Copy Data to Clipboard”
11) Paste into Excel or other spreadsheet software
12) Find the “Response” column

a) This will tell you the extent of binding at each concentration.
13) Calculate the % binding for each competitive analyte step where step 10 (following primary analyte blank) is 100%.

a) Competing VHHs will have a flat line while non-competing VHHs will show a second increase (**Figure 20A**).
b) Competing VHHs will show differing amounts of response depending on the saturation level of the primary VHH (**Figure 20B**).
14) Prepare an XY plot

a) X = Concentration of primary analyte
b) Y = Competitive analyte % of max binding
15) Fit a dose-response curve (**Figure 20C**)
16) Repeat experiment with primary and competitive analytes swapped.

**Figure 18:**
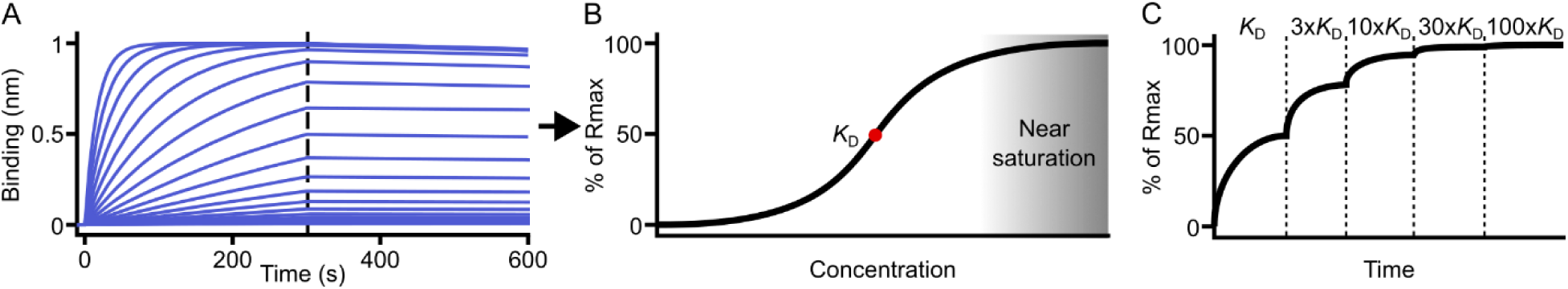
relationship between saturation and concentration. A) Samples reach maximal binding depending on the sensor capacity at a rate determined by kinetic parameters and concentration. B) [*K*_D_] always results in 50% saturation at steady-state, and increasing doses asymptotically approach saturation. C) Estimation of sufficient saturation for a competition assay can be measured with back-to-back association steps with escalating concentrations and will plateau at higher concentrations. This plot allows each concentration to reach steady-state, but a slow-forming complex may require much greater than 100×*K*_D_ to reach near-R_max_ in a reasonable timeframe.

**Figure 19:**
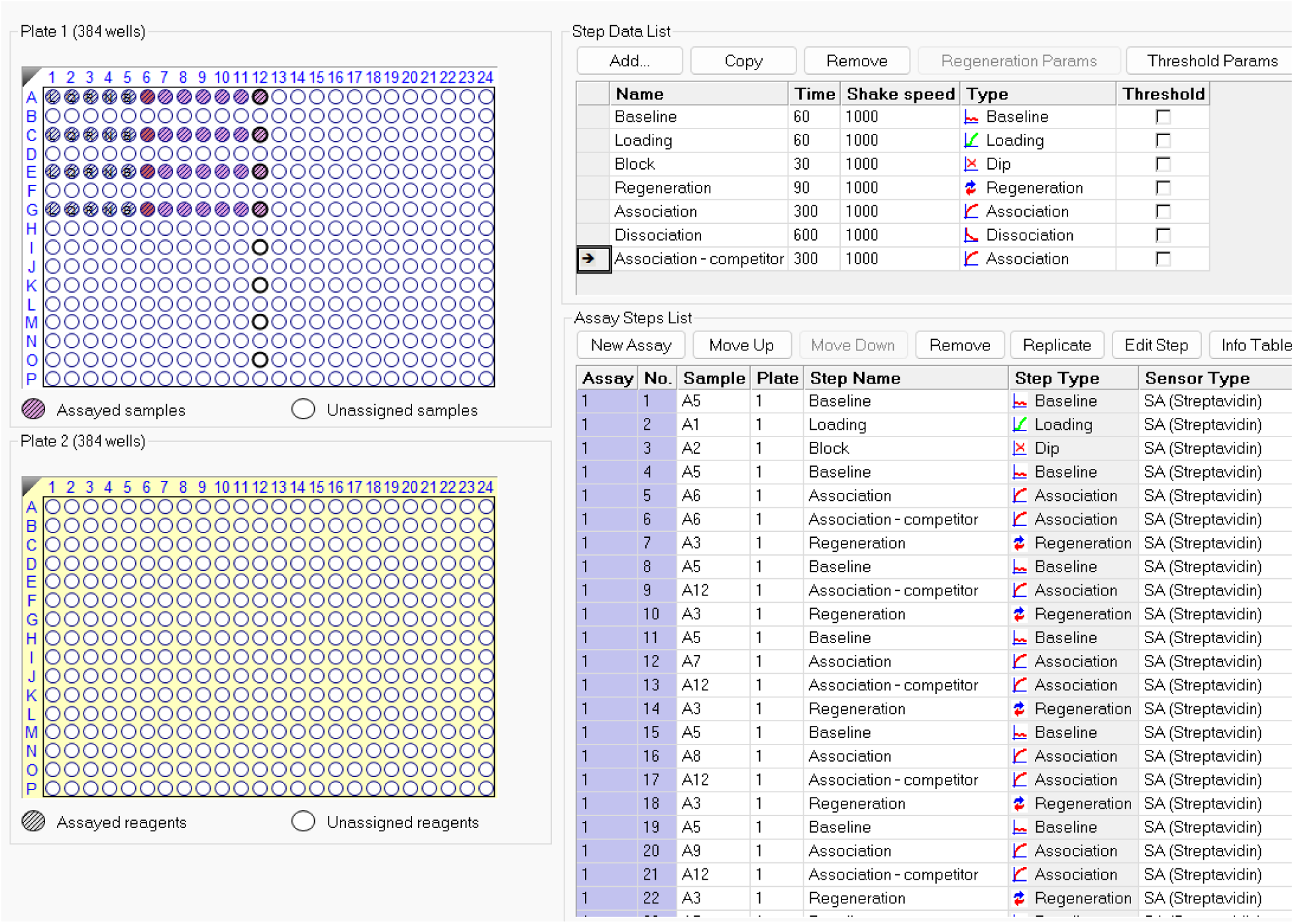
Sample assay design for a competition experiment. See the attached importable method file and supplemental methods spreadsheet for additional details

**Figure 20:**
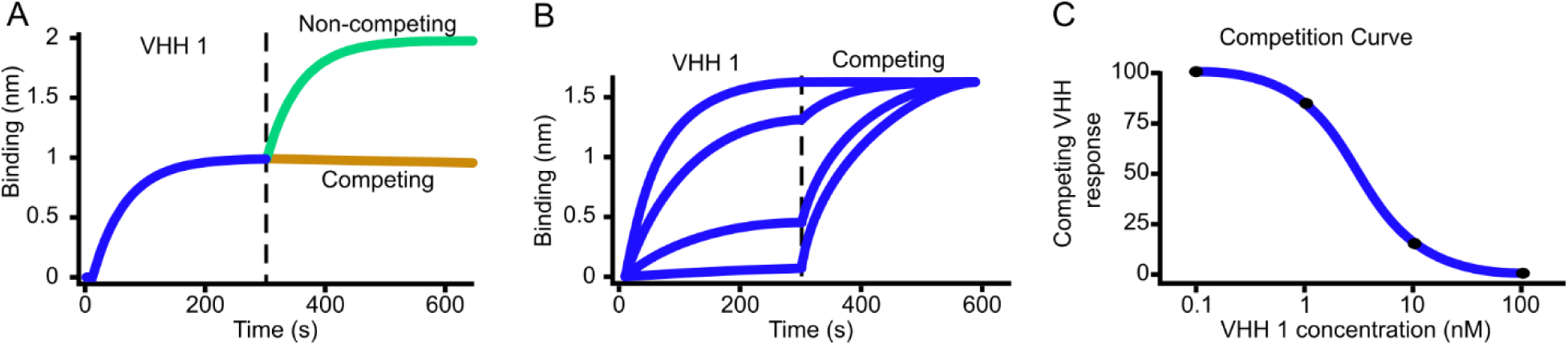
Interpretation of competition results. (A) Sensorgram of VHHs showing competition or non-competition with VHH1, assuming that both bind to the target ligand. (B) Dilution series of VHH 1 reaching various levels of saturation partially block binding of the competing VHH. If the competing VHHs are not the same size, then all concentration combinations will not tend toward the same final response value. (C) dose-response curve showing decreasing competing VHH response with logarithmically increasing VHH1 concentration.

### Procedure 4: Epitope binning

Timing 3 hours

1. Perform pilot experiments to determine the saturating concentrations for all of the different analytes to be tested.
2. Start a new method using the epitope binning template in the wizard or start with a previous epitope binning method (see example method file in the Supplemental Information).
3. Set up the sample plate according to the example in (**Figure 21**).
4. The key to this experiment is to have the first sample column contain each different analyte in separate wells. Up to 8 analytes per experiment can be tested this way, if using an 8 channel Octet. Subsequent columns should contain just one analyte in every row of the column. Thus by treating column 1 as the primary analyte and stepping through the competitive analyte rows, every analyte will be tested against every other, including itself.
5. Set up the step times similarly to Procedure 3 (see example plate setup in **Figure 22**).

a. Note that the competitor analyte association step is considerably shorter than the primary analyte saturation step.
6. Run as normal, following Procedure 1 steps 9-16.
7. To analyze, open the file in Data Analysis HT
8. Click the “Epitope” button in the top bar
9. If designed as above, the software should automatically recognize the binding cycles and generate a matrix. **Figure 23** shows an example of this with a simple binning experiment.
10. This experiment can be repeated to include more sample analytes, or to include analytes with known binding sites.

a. If 16 analytes require testing with an 8 channel machine, experiment 1 will load analytes 1-8 on tips and test against all 16 analytes whereas experiment 2 will load analytes 9-16 and again test against all 16. This method can be extended to cover any number of analytes using only 1 tip per additional analyte, as long as the total experiment time does not exceed the 3 hour limit due to evaporation.
11. This method is suitable for high-throughput screening depending on the type of Octet machine available (up to 96 channels), but key results should be validated with the quantitative competition experiment to verify that blocking is dose-dependent.
12. Because this method does not use replicate tips, it is important to repeat the entire experiment at least twice.

**Figure 21:**
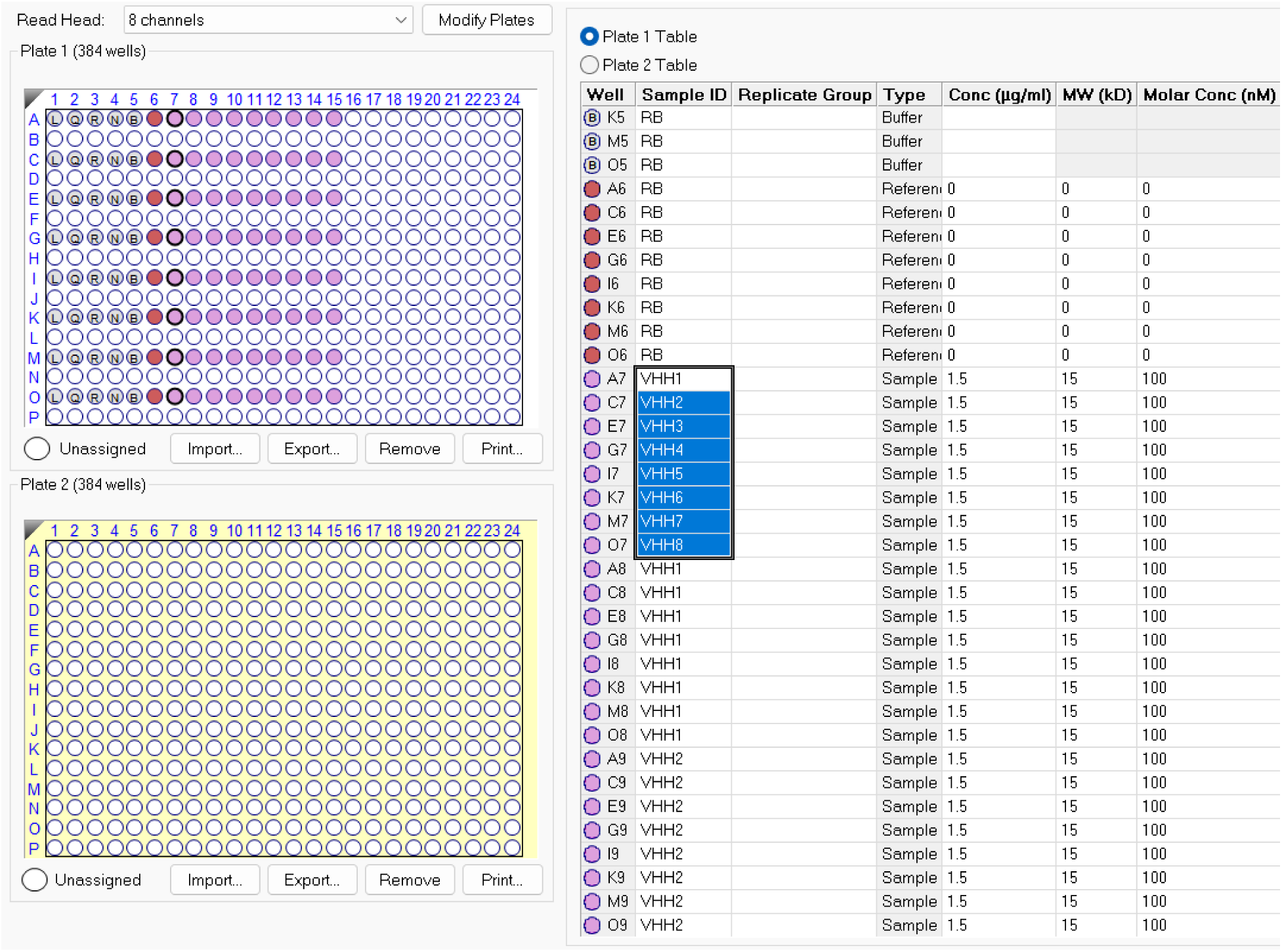
Example plate layout for epitope binning experiments.

**Figure 22:**
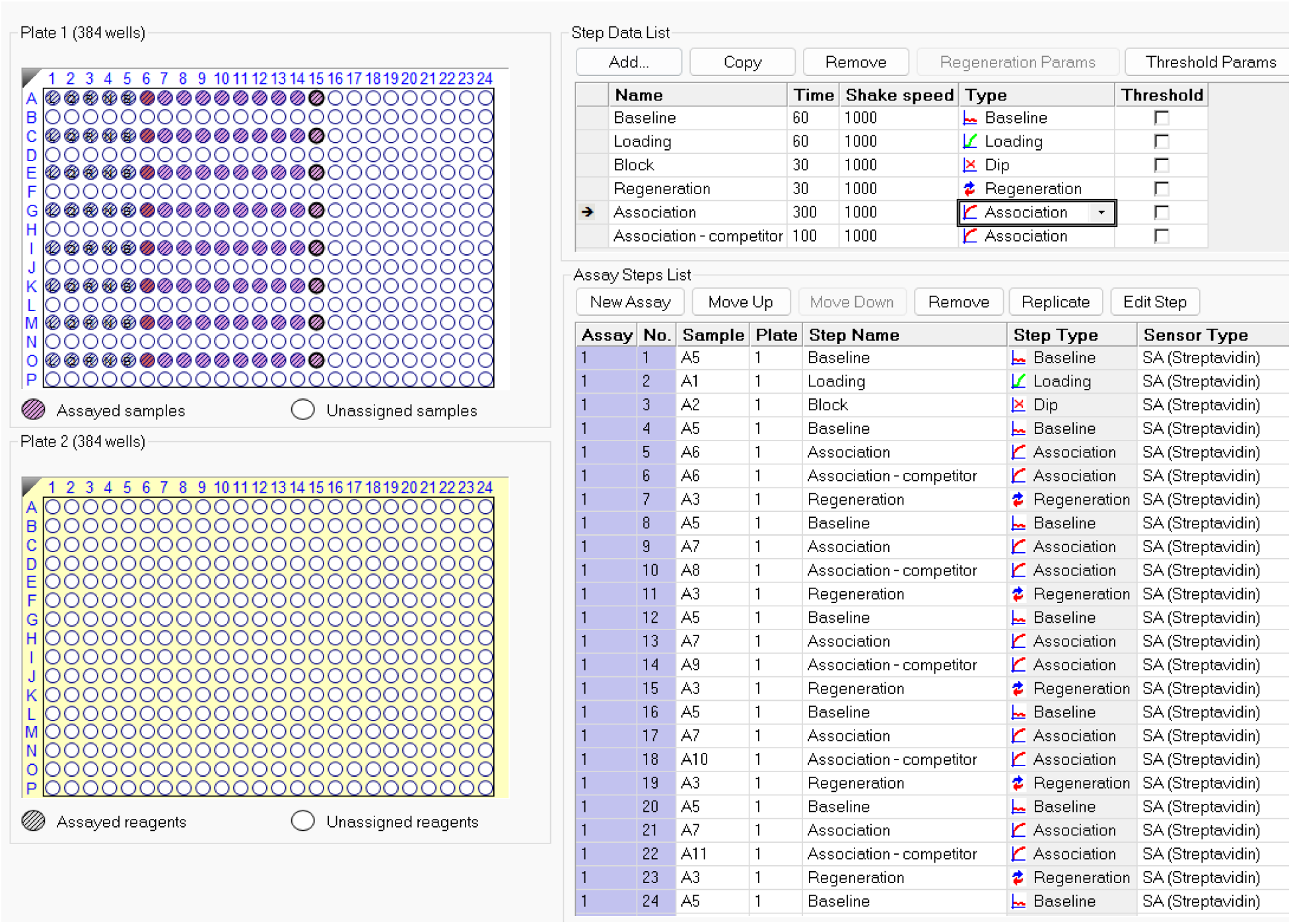
Example step order and timing for epitope binning experiments.

**Figure 23:**
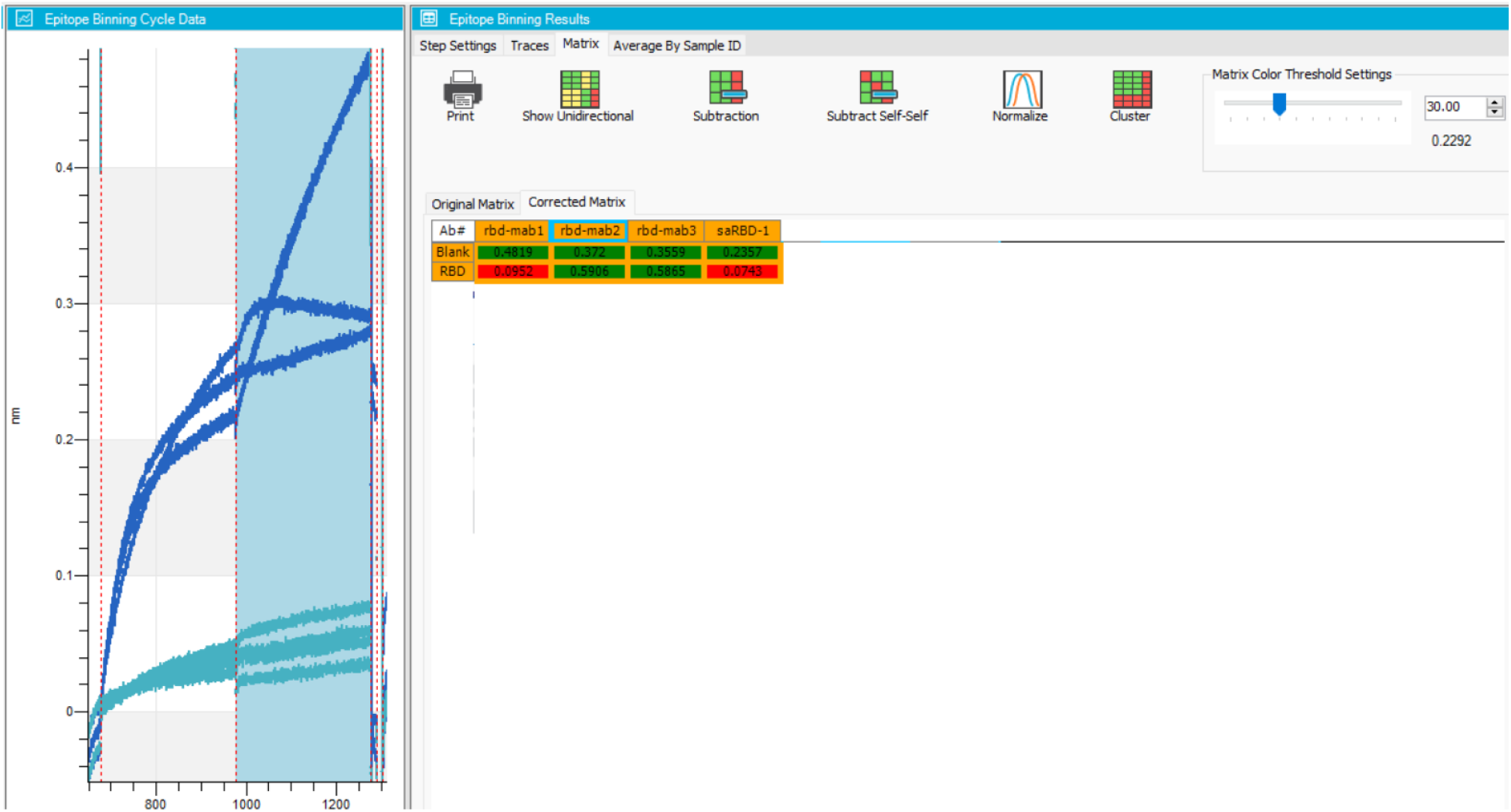
Epitope binning analysis example. In this example, saRBD-1 was used as the primary analyte for all cycles followed by 3 different monoclonal antibodies or saRBD-1 again. We can see that saRBD-1 competes with itself, a good positive control that saturation worked correctly, and mab1 but not mab2 or mab3 competes with saRBD-1. This indicates that saRBD-1 likely binds to the same site as mab1, a known class 1 Spike-binding antibody.

### Optional Procedure 1: Surface biotinylation of target protein

The recommended method for preparing biotinylated ligands for initial BLI testing is succinimide-based Lysine labeling. It is preferred to use a method which allows for quantification of biotin incorporation following labeling. For this purpose, we recommend ChromaLINK Biotin (Vector Labs, Cat no. B-1001). For convenience (at the expense of cost) this reagent is also available as a kit containing other essential items (Vector Labs, Cat no. B-9007-105K) such as DMF, modification buffer, Zeba columns, and control samples. Knowledge of the protein’s amino acid sequence is important for this process to verify the number of available lysines and to determine the extinction coefficient (https://web.expasy.org/protparam/). Proteins without lysines may still be labeled at the N-terminus. Biotinylation can be performed using the manufacturer’s user guide or using the following protocol which prepares 100µg of labeled protein. This is sufficient for 10-20 BLI experiments with 8-16 tips each.

#### Materials

1. Protein of interest, ideally 100 µL at approximately 1 mg/mL

- If possible, prepare without amine-containing buffer salts (Tris, glycine, etc.)
2. ChromaLINK reagent (Vector Labs, Cat no. B-1001)
3. Anhydrous DMF (Vector Labs, Cat no. S-4001-005)
4. 1 - Zeba spin column (Fisher Scientific, Cat no. PI89882)
5. 1 mL modification buffer (100mM Na_2_HPO4, 150mM NaCl, pH 8.0)
6. 1 mL Final buffer (See main protocol, typically 1x PBS)

#### Equipment

1. Microcentrifuge (Fisher Scientific, Cat no. 13-100-675)
2. Micropipettes (0.1–2.5, 2–20, 20–200 and 100–1,000 µL; Eppendorf, Cat. nos. 3123000012, 3123000098, 3123000055 and 3123000063)
3. Micropipette refill tips (10, 200 and 1,000 µL; STARLAB, Cat. nos. S1111-3700, S1111-0706-C and S1111-6701-C)
4. 1.5 mL Microcentrifuge tubes (Greiner, Cat no. 616281)
5. Nanodrop (Thermo Scientific, 840-329700)

#### Software

1. ChromaLINK Biotin Conjugation Calculator (find attached or at the following link)

- https://vectorlabs.com/productattachments/protocol/VL_B-9007-105_Calculator_LBL02107.xlsx

#### Procedure

1. Buffer exchange protein into modification buffer

a. Wash Zeba column 3× with 500 µL modification buffer

i. Spin 1 min at 1,500×g at room temperature
b. Add no more than 120 µL of target protein solution

i. Capture eluted protein in a fresh 1.5mL microcentrifuge tube
ii. Spin 2 min at 1,500×g at room temperature
iii. Immediately add 500µL DI water to the used Zeba column
2. Determine concentration of desalted protein

a. Measure the extinction coefficient corrected OD_280_ by Nanodrop
3. Prepare the ChromaLINK reagent

a. Dissolve 1 fresh tube of reagent to 5 mg/mL in anhydrous DMF

i. Keep tube closed tightly when not in use
ii. Store at −80°C
iii. Let warm to room temperature before opening to avoid condensation
4. Use the ChromaLINK Calculator to determine appropriate reagent ratio

a. 5-10 equivalents is typically suitable to achieve labeling with 1-2 biotin/protein
5. Add the indicated quantity of ChromaLINK stock

a. Mix well by pipetting
6. Incubate at room temperature, away from light, for 1-2 hours.

a. Solution will likely become cloudy, this does not necessarily indicate protein loss.
7. Meanwhile, wash the Zeba column used in step 1

a. 3× 500µL DI water, 1,500×g spin for 1 min at room temperature
b. 3× 500µL final buffer, 1,500×g spin for 1 min at room temperature
c. Wait until step 6 completes for the final spin to avoid drying beads.
d. Alternatively use a fresh Zeba column and wash 3× final buffer.

i. Contamination with small quantities of unlabeled protein is not a concern for downstream steps, so a fresh column is unnecessary.
8. Desalt labeling reaction to remove free probe*

a. Add full reaction volume (<130µL) to equilibrated the Zeba column

i. Spin 1,500×g for 1 min at room temperature
9. Measure labeling efficiency

a. Measure OD_280_ and OD_354_ values in manual mode by Nanodrop

i. Ensure values are corrected for 1-cm path length
ii. Expected recovery is over 95% for stable proteins
iii. Reasonable OD_280_ values will be 0.5-2.0 depending on ext. coef.
iv. Reasonable OD_354_ values will be ∼5-30% higher than OD_280_.
b. Use the E1% tab of the calculator to determine biotins/protein labeling
10. Store protein

a. Aliquot protein (5-10µL per tube)
b. Snap freeze and store at −80°C until needed.
c. Also freeze any aliquots to be used immediately to ensure consistency between samples, or specifically test fresh versus frozen.
d. Properly record information about input protein, labeling method, and labeling efficiency. Update with information about BLI tip loading efficiency after the first pilot experiment is complete.

*Some detectable free probe will remain after this desalting method (Zeba). This will result in slightly reduced apparent capacity of the biosensors as free probe competes for streptavidin on the sensor. This is not typically problematic, but more complete desalting can be achieved using dialysis if preferred.

### Timing

Antigen preparation using the biotinylation kit: 3 hours

Pilot experiment: 3 hours

Follow up experiment using procedures 2, 3 or 4: 3 hours each

### Troubleshooting

Troubleshooting information can be found in Table 5

**Table 5:** Troubleshooting information.

| Procedure | Step | Problem | Possible reason | Solution |
| --- | --- | --- | --- | --- |
| 1 | 2 | Method file cannot be opened | Wrong instrument setting | Go to File>options and verify that the correct instrument type is selected under "simulation." |
| 1 | 11 | Sudden shifts at random times during steps | Sensor damage | Ensure proper handling of sensors, do not touch tips to any surface. Avoid physical jolts or moisture in storage. |
| 1 | 14 | Poor loading | Failed biotinylation | Check biotin incorporation and repeat as necessary. Verify pH of reaction buffer. |
| 1 | 14 | Poor loading | Free biotin contamination | Repeat buffer exchange using desalting or dialysis |
| 1 | 14 | Poor binding | Inactive ligand | Verify ligand purity and quality by SDS-PAGE, Western Blot, ELISA, and/or functional assay. Repurify by size exclusion, or try alternative constructs or sources. Also try alternative tagging/labeling and sensor chemistry. |
| 1 | 14 | Poor binding | Impure analyte | Verify analyte purity and quality by SDS-PAGE, Western Blot, ELISA, and/or functional assay. Repurify by size exclusion, or try alternative constructs or sources. Also try alternative tagging/labeling and sensor chemistry. |
| 1 | 14 | Poor binding | Low analyte concentration | Raise maximum analyte concentration. Ensure proper background controls. |
| 1 | 14 | Poor binding | Surface interaction artifact | Swap ligand and analyte. |
| 1 | 14 | Poor binding | Small analyte | Generate a larger analyte construct or try a competition experiment. |
| 1 | 14 | Negative binding signal | Large particles | Excessively sized particles can produce negative signal upon binding. Data can be inverted for quantitation experiments only as of Analysis HT version 10.0. |
| 1 | 14 | High background | Sticky analyte | Change blocking/running buffer |
| 1 | 14 | High background | Sticky control | Change loading control ligand |
| 1 | 14 | Consistent drift up or down throughout run | Normal sensor drift | This is not concerning as long as double reference subtraction results in a flat baseline. |
| 1 | 14 | Steep downward drift | Ligand unbinding | Use a control sensor for drift, or change tip chemistry. |
| 1 | 14 | Escalating upward drift during baseline | Buffer well contaminated with ligand | Change buffer well for each cycle. |
| 1 | 14 | Steep upward drift | Sticky ligand | Modify running buffer or use each sensor for fewer cycles. |
| 1 | 14 | Baseline is flat, but starting value increases with each cycle. | Incomplete regeneration | Increase strength of regeneration buffer or try different chemistry. Not all ligands can be fully regenerated, use only one cycle per tip until suitable regeneration can be established. Try reloadable tips like anti-His and reload with fresh ligand at each step. |
| 1 | 14 | Loss of signal over cycles | Unstable ligand | Not all ligands can be regenerated, use only one cycle per tip until suitable regeneration can be established. Try reloadable tips like anti-His and reload with fresh ligand at each step. |
| 1 | 14 | Rapid binding (>5 sec) followed by slower, normal signal development. | Extreme buffer change | Verify buffer composition of analyte stock. Buffer exchange more completely or concentrate stock to allow greater dilution in running buffer. |
| 1 | 14 | Sharp spikes of >5nm | Air bubble | Ensure proper liquid level in wells and no trapped air bubbles. |
| 1 | 14 | Sharp spikes of >5nm | Extreme buffer change | Verify buffer compositions. |
| 1 | 25 | $R_{max}$ decreases after regeneration(s) | Ligand damage due to incompatible regeneration | Try different regeneration times/buffers. Use each sensor for only 1 cycle. Do not attempt repeated single-cycle kinetics until a suitable analysis method has been established and tested. |
| 1 | 38 | Binding too fast to see curvature | Excessively fast on-rate | Use steady-state kinetics. Verify that shift is not due to buffer components. |
| 1 | 38 | No signal drop during dissociation | Excessively low off-rate | Extend dissociation step. If less than 5% signal drop after 5000 seconds, report $k_{OFF} < 1 \times 10^{-5} \text{ s}^{-1}$. |
| 1 | 38 | Two-phase dissociation or association | Non-specific binding | Change blocking/running buffer. |
| 1 | 38 | Two-phase dissociation or association | Non-langmuirian kinetics | Test for non-standard binding in other assays. Possible analyte-induced conformational change. |
| 1 | 38 | Odd curve shape | Non-specific binding | Check control tips, use alternative loading control, change blocking/running buffer. |
| 1 | 38 | Odd curve shape | Non-langmuirian kinetics | This is evidence that the 1:1 model does not explain the observed binding. Test stoichiometry of binding using alternative methods. |
| 1 | 38 | Poor fit | Outlier concentrations | Exclude highest and/or lowest concentrations from calculations. Unusual curves at higher concentrations may be caused by non-specific binding. |
| 2 | 13 | Poor fit | Non-langmuirian kinetics | This is evidence that the 1:1 model does not explain the observed binding. Test stoichiometry of binding using alternative methods. |

### Anticipated results

BLI experiments provide binding data regarding a selected interacting pair, but this can be either quantitative measurement of binding rate constants or qualitative information about overlapping epitopes. Competition experiments can also be designed to test the binding of molecules which would otherwise be too small to detect using standard BLI setups. Further, epitopes and biological functions can be interrogated by creating panels of mutants or naturally occurring variants. Examples of these techniques being used to answer biological questions can be found in our studies^8–15^, and countless others are not only studying novel biology but creating new BLI-based methods^27,42–45^.

Proper interpretation of BLI data is as important as correctly setting up the experiment. When communicating BLI data it is essential to provide enough information about any biomolecules used including sequence, purity, concentration, buffer composition, and labeling methods. Any information about multimer formation under different conditions can also help with interpreting results, but BLI can often be used to identify whether interactions other than simple 1:1 binding are occuring. **Figure 24A** shows raw, double-referenced binding curves for the self-association of the Tuberculosis protein ESAT-6 at pH 4.5. These curves do not fit well to a single exponential because of the formation of larger molecular weight oligomers, which also make regeneration less efficient through formation of large, stable complexes. However, it is not always possible to tell from the data that a multivalent interaction is occurring. Traditional antibodies or bivalent nanobody constructs are a good example. **Figure 24B** shows raw, double-referenced binding curves for bivalent saRBD-1 which appears to fit reasonably well to a single exponential.

**Figure 24:**
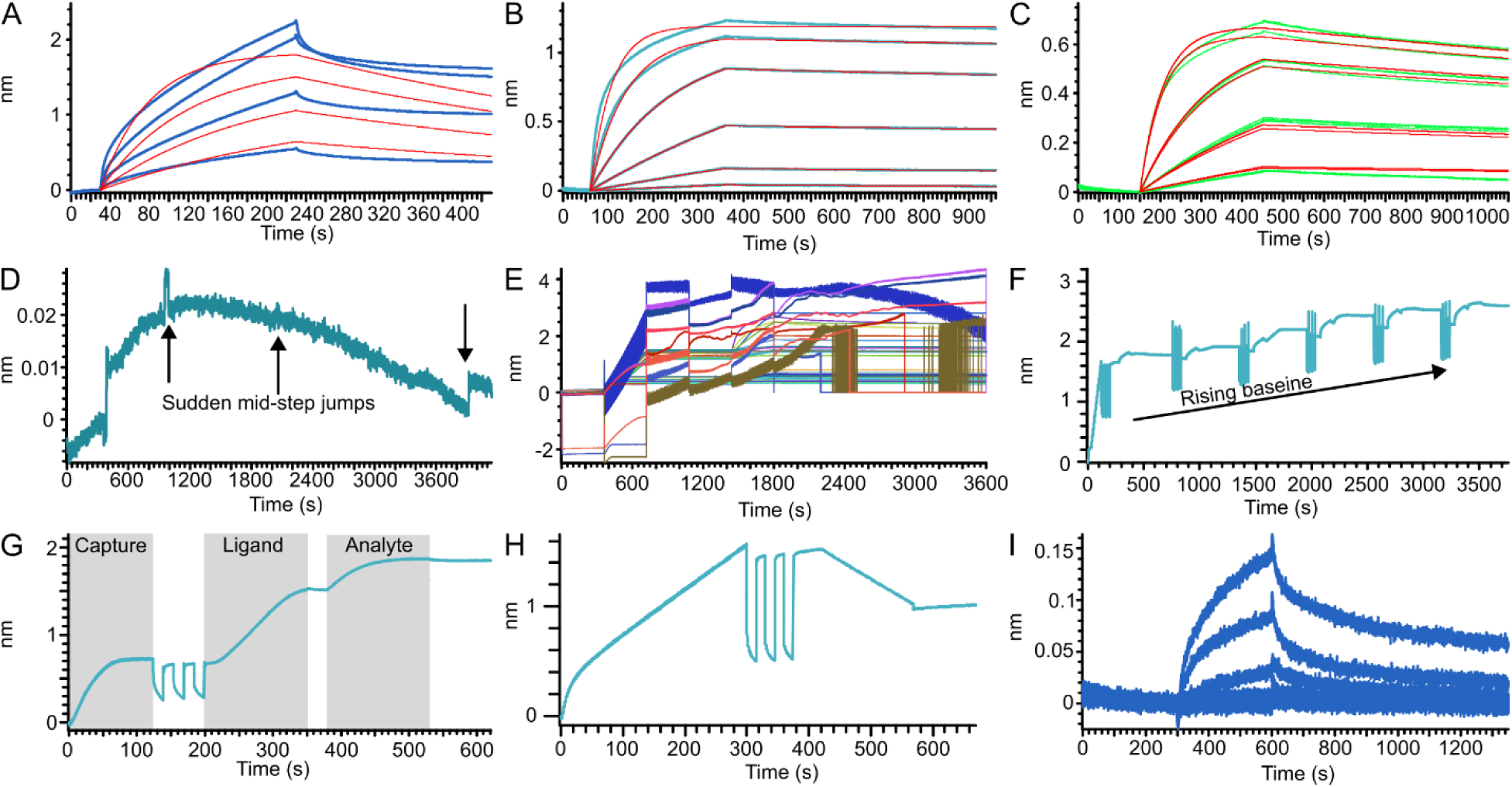
Example data showing common results. Sensorgrams showing: (A) complex binding not fitting a 1:1 model due to formation of large multimers. (B) A bivalent nanobody complex displaying acceptable fitting to a 1:1 model despite not being a 1:1 interaction. (C) A true 1:1 interaction run in triplicate and globally fit. (D) A damaged sensor tip showing signal jumps at irregular intervals. (E) Insufficient liquid volume due to excess evaporation causing widespread artifacts. (F) Rising baseline over multiple binding cycles due to incomplete regeneration. Also note the decreasing R_max_ with each new cycle. (G) A sandwich experiment using a biotinylated capture nanobody which is subsequently loaded with ligand (antigen) and measured with analyte (a different nanobody). Regeneration will remove everything except the capture nanobody, allowing fresh antigen to be captured for later rounds. (H) Negative binding signal from a large (∼30 nm) nanoparticle. (I) Low signal-to-noise due to poor analyte binding.

This is why proper reporting of materials is key to successful interpretation. It is possible to measure a proper *K*_D_ for a (bivalent) monoclonal antibody, but the orientation must be such that the analyte cannot have multiple points of contact with the sensor surface. This can involve digestion of antibodies to make Fabs, but if the antigen only contains a single binding site and does not multimerize then the antibody can be used as the ligand and the antigen as analyte. It may also be possible to attach the antigen at a low enough density to the sensor that antibodies are not able to bridge two epitopes, but this would need to be done with suitable care.

When troubleshooting a new BLI experiment, it is important to be able to recognize successful and unsuccessful experiments. **Figure 24C** shows a high-quality experiment in triplicate that fits well with a global 1:1 model. Various major issues can be identified by looking at the raw sensorgram traces such as biosensor damage (**Figure 24D**) or insufficient volume (**Figure 24E**). If only one well is showing the issues seen in **Figure 24E**, then there may be a bubble in that well. It is also possible to see more subtle errors such as insufficient regeneration/washing (**Figure 24F**). Ideally, the baseline of each cycle should start from approximately the same nm value. Decreasing signal may indicate ligand falling off of the sensor. This can be addressed by re-loading with ligand, however whether this is possible will depend on sensor chemistry. Biotin should not measurably dissociate over the timeframe of an experiment. Another option with sensitive or multi-component ligands is to use a sandwich approach in which a known nanobody/antibody is attached to the sensor and then fresh loading steps can be used for each cycle to provide a consistent amount of fresh ligand (**Figure 24G**). In contrast, a signal which looks normal, but inverted may be due to extremely large particles such as liposomes or nanoparticles (**Figure 24H**). One explanation of this is that inefficient packing of crowded large particles creates a rarified area proximal to the sensor surface.

Weak binding or insufficient loading is also identifiable by a reduction in signal-to-noise ratio (**Figure 24I**) which is observable as an apparent broadening of the traces. There is a fairly consistent amount of natural sub-second variation which can be used to assess signal-to-noise. Noise is typically 0.02-0.03 nm using default settings (before Savitzky–Golay filtering) and using a tilted-well 384-well plate. Low signal can be addressed by attaching more ligand, verifying ligand/analyte quality, and ensuring that ligand is free of contamination from interfering compounds like free biotin. Some analytes are simply too small or too weakly binding to give sufficient signal in BLI.

Interpretation of competition experiments is simpler because quantification is not necessary. It must be verified through pilot studies that the interaction of ligand with all analytes is consistently working, and partial phenotypes may indicate that concentrations are not saturating. Determination of orthosteric versus allosteric competition can be challenging. While this may be partially addressed with BLI using a conformational change model (not currently implemented in Octet software), other confirmatory approaches must also be used (allosteric paper, Harris, mighty).

When presenting BLI data, authors should report experimental design including concentrations, step times, temperature, and sensor types. For each experiment, authors should show raw curves with fit lines overlaid for each concentration tested and fitting parameters including model (this should always be 1:1), and fitting method (i.e. global, Rmax unlinked by sesnor) should be provided. Where an interaction is not 1:1 and it is not possible to adjust the experimental design, it is often best to use a 1:1 model anyway and clearly report the value as an apparent *K*_D_ rather than a true *K*_D_. This can at least provide an objective point of comparison for other work.

It is often not possible to show all replicates in individual figures without crowding the plots, but the number of replicate sensors or experiments should be stated and the method of combining replicate data should be explained. If *K*_D_ values are reported, then errors should be provided, with the understanding that the error provided by the software is only the fitting error and does not describe experimental error that can only be found by comparing multiple distinct experiments. Dr. Rebecca Rich and Dr. David Myszka wrote an excellent series of critical reviews highlighting best practices and poor examples of kinetic binding data in published literature every year between 1998 and 2009 ^22,46,47^. While biosensor technology has improved steadily in the time since these were published, the lessons contained are timeless and yet to be fully implemented.

## Supporting information

Example method files

## Author contributions

T.A.B. Wrote the manuscript and designed the protocols. S.K.G., J.B.W., M.TG., A.A., T.A., A.H., and X.N. assisted with assay development. U.S., J.E.B., and F.G.T. Supervised the research. All authors reviewed and edited the manuscript.

## Acknowledgements

We thank the Tafesse, Shinde, and Burke labs for their collaboration and support. T.A.B. is supported by NHLBI training grant 5T32HL083808. J.E.B. is supported by the Canadian Institutes of Health Research (CIHR, 168998), and the Michael Smith Foundation for Health Research (MSFHR, scholar 17686). FGT is supported by NIH grant R01AI141549, and the OHSU Silver Innovation Award. BLI data were generated on an Octet Red 384, which is made available and supported by the OHSU Biophysics Shared Resources Core and equipment grant number S10OD023413.

## Competing interests

J.E.B. reports personal fees from Scorpion Therapeutics, Reactive therapeutics and Olema Oncology; and research grants from Novartis. All other authors declare that they have no competing interests.

## Data and Code Availability

The datasets generated during and/or analysed during the current study are available from the corresponding author on reasonable request. No unique code was generated as part of this work.

